# An AAV capsid reprogrammed to bind human Transferrin Receptor mediates brain-wide gene delivery

**DOI:** 10.1101/2023.12.20.572615

**Authors:** Qin Huang, Ken Y. Chan, Shan Lou, Casey Keyes, Jason Wu, Nuria R Botticello-Romero, Qingxia Zheng, Jencilin Johnston, Allan Mills, Pamela P. Brauer, Gabrielle Clouse, Simon Pacouret, John W. Harvey, Thomas Beddow, Jenna K. Hurley, Isabelle G. Tobey, Megan Powell, Albert T. Chen, Andrew J. Barry, Fatma-Elzahraa Eid, Yujia A. Chan, Benjamin E. Deverman

**Affiliations:** Stanley Center for Psychiatric Research, Broad Institute of MIT and Harvard; Cambridge, USA; Department of Systems and Computer Engineering, Faculty of Engineering, Al-Azhar University; Cairo, Egypt

## Abstract

Developing vehicles that efficiently deliver genes throughout the human central nervous system (CNS) will broaden the range of treatable genetic diseases. We engineered an AAV capsid, BI-hTFR1, that binds human Transferrin Receptor (TfR1), a protein expressed on the blood-brain barrier (BBB). BI-hTFR1 was actively transported across a human brain endothelial cell layer and, relative to AAV9, provided 40–50 times greater reporter expression in the CNS of human *TFRC* knock-in mice. The enhanced tropism was CNS-specific and absent in wild type mice. When used to deliver *GBA1*, mutations of which cause Gaucher disease and are linked to Parkinson’s disease, BI-hTFR1 substantially increased brain and cerebrospinal fluid glucocerebrosidase activity compared to AAV9. These findings establish BI-hTFR1 as a promising vector for human CNS gene therapy.

## Introduction

A critical challenge in the development of more efficient delivery vectors for gene therapy is engineering vectors with known mechanisms of action (MOAs) relevant to human patients. Conventional approaches to engineering vectors such as adeno-associated viruses (AAVs) with enhanced tropisms have largely relied on capsid library selections in animals. These selections have been used by numerous groups over the past two decades because they can be successful without requiring prior knowledge of an MOA (*1–7*). However, vectors selected in animals mostly have not translated across preclinical models. Despite extensive searches, AAV capsids with clear translational potential and known MOAs for entering the CNS have not yet been described. Here, we approached this challenge from a different perspective and selected AAV capsids first for a specific MOA, that is binding to the human Transferrin Receptor (TfR1), and showed that one of these capsids crossed the blood-brain barrier (BBB) and mediated efficient gene delivery throughout the CNS.

We chose TfR1 as a target because of its high expression on the human BBB; its ability to mediate constitutive, ligand-independent receptor-mediated transcytosis (RMT) across the CNS vasculature (*8–12*); and its track record as a target to increase the delivery of biologics into the CNS of mice (*13–15*), nonhuman primates (NHPs) (*16–18*), and humans as investigational therapies (*19–22*) and as an approved antibody-based therapeutic for Mucopolysaccharidosis type II (*23*). We first screened 7-mer-modified AAV9 capsid libraries for their ability to bind to human TfR1 *in vitro*. The top-performing capsid, AAV-BI-hTFR1, exhibited more efficient gene delivery to human brain endothelial cells and improved active transport across a human vascular monolayer. When systemically administered to adult *TFRC* knock-in (KI) mice carrying a chimeric *Tfrc* gene with a humanized extracellular domain, BI-hTFR1 transduced the majority of neurons and astrocytes across multiple brain regions. The enhanced tropism was entirely dependent on an interaction with the humanized TfR1, as we observed no enhancement in wild type (WT) mice. The tropism was also selectively enhanced in the CNS, consistent with the high level of expression of *TFRC* on the CNS vasculature relative to the vasculature in other organs (*12*).

To explore its ability to deliver a therapeutically relevant cargo, we intravenously injected *TFRC* KI mice with BI-hTFR1 or AAV9 packaging the human Glucosylceramidase Beta 1 (*GBA1*) gene. Inactivating mutations of *GBA1* are the primary cause of Gaucher’s disease, a lysosomal storage disorder often affecting the CNS, and *GBA1* mutations are a genetic risk factor for Parkinson’s disease and dementia with Lewy Bodies (*24*). Systemic delivery of BI-hTFR1:*GBA1* in adult *TFRC* KI mice resulted in increased expression throughout the brain and elevated glucocerebrosidase (GCase) activity in the brain and cerebrospinal fluid (CSF), which were not observed with AAV9. Through its direct engagement of TfR1, a receptor of natural protein ligands as well as therapeutic biologics in humans, BI-hTFR1 represents a promising vector for the development of CNS-targeting human gene therapies.

## Results

### AAV capsids engineered to bind human TfR1

To target AAV capsids to TfR1, we screened AAV9-based NNK capsid libraries of variants with random 7-mer insertions between VP1 residues 588–589 for selective binding to human TfR1 using our recently described receptor-targeting approach (*25*) (Fig. 1A). Based on the results of the *in vitro* screens, we chose four capsids that share a common sequence motif for validation: BI-hTFR1, BI-hTFR1-2, BI-hTFR1-3, and BI-hTFR1-4. We used each capsid to individually package a single-stranded genome that expresses nuclear mScarlet and Luciferase under the control of the ubiquitous CAG promoter (ssAAV-CAG-NLS-mScarlet-P2A-Luciferase-WPRE-SV40pA). The four capsids exhibited elevated association with and transduction of CHO cells that stably express human *TFRC* but not the control CHO cells or those that express *TFRC* from rhesus macaque, marmoset, or mouse (Fig. 1B and C, and fig. S1). We next tested whether the capsids were capable of increased association with and transduction of human brain endothelial cells that endogenously express *TFRC*. Although the extent of AAV associated with primary human brain microvascular endothelial cells (hBMVECs) and a human brain endothelial cell line (hCMEC/D3) varied (Fig. 1D), each of the four TfR1-binding capsids exhibited significantly enhanced transduction of both brain endothelial cell types compared to AAV9 (Fig. 1E). To further assess the relative binding affinities of the capsids with purified human TfR1, we performed bio-layer interferometry (BLI). We derived kinetic constants using a multi-phasic binding model because a monovalent interaction algorithm led to a poor fit of the data. Factors that may contribute to the multi-phasic binding curves are outlined in the Materials and Methods section. Based on its performance across cell association and transduction assays and its ability to bind human TfR1 at lower concentrations (Fig. 1F and fig. S2A), we chose BI-hTFR1 for further investigation in the rest of this study.

**Fig. 1.**
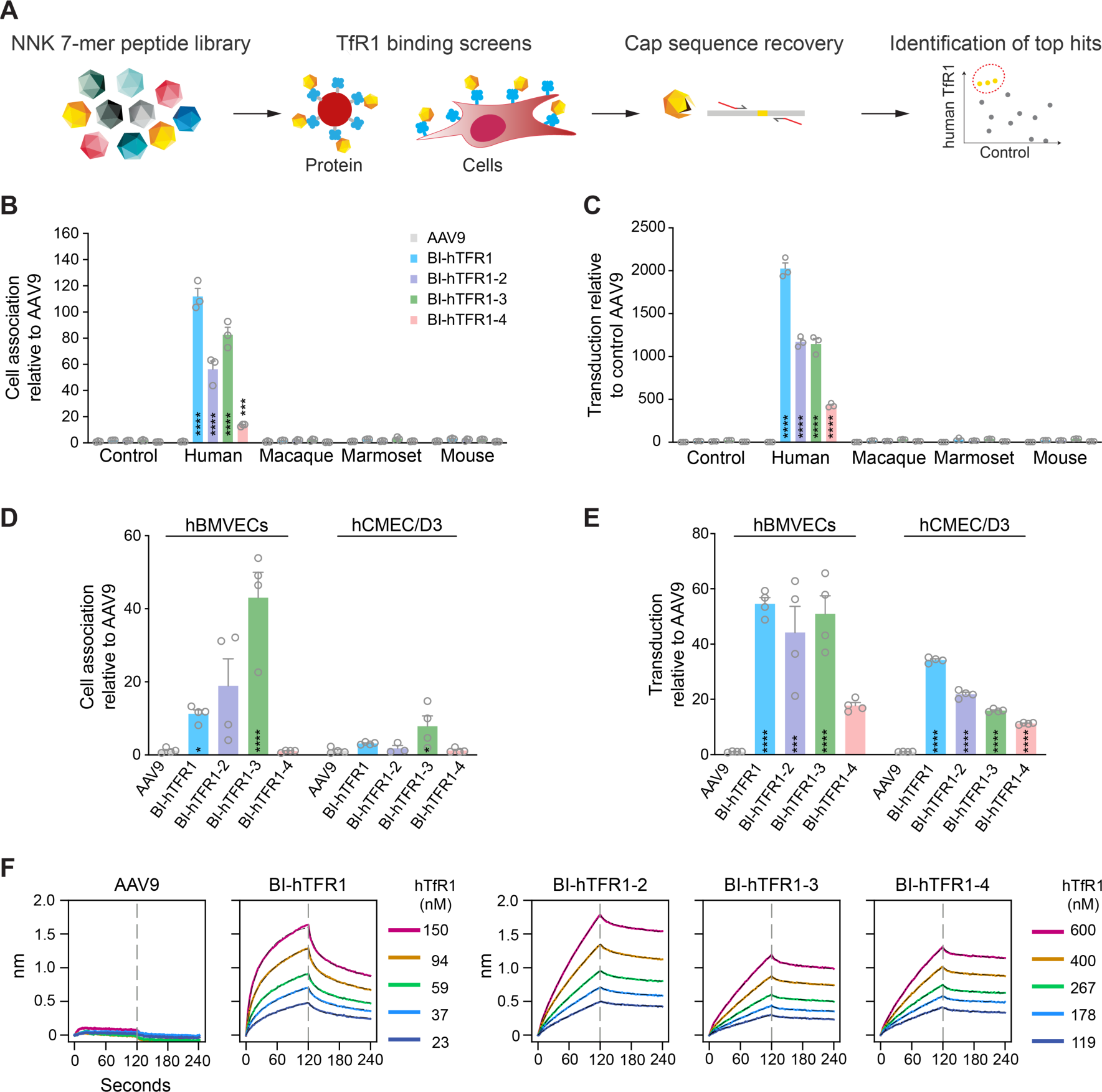
AAV9 can be programmed to bind human TfR1. (**A**) An AAV9-based NNK capsid library of variants with random 7-mer insertions between VP1 residues 588–589 was screened for selective binding to human TfR1 in pull-down or cell binding assays. Individually produced human TfR1-binding variants carrying a CAG-NLS-mScarlet-P2A-Luciferase-WPRE-SV40pA construct exhibited enhanced species-specific (**B**) association with and (**C**) transduction (Luciferase activity) of CHO cells stably expressing *TFRC* (two-way ANOVA using AAV9 as the main comparison group for each cell line with Bonferroni multiple comparison correction: **** and *** indicate *p* ≤ 0.0001 and ≤ 0.001, respectively; *n* = 3 replicates, error bars indicate ± SEM). Values reported are normalized to AAV9 in each cell line. The TfR1-binding variants exhibited enhanced (**D**) association with and (**E**) transduction (Luciferase activity) of hBMVEC and hCMEC/D3 cells (one-way ANOVA using AAV9 as the comparison group for each cell line with Dunnett’s multiple comparison correction: ****, ***, and * indicate *p* ≤ 0.0001, ≤ 0.001, and ≤ 0.05, respectively; *n* = 4 replicates, error bars indicate ± SEM). Values are normalized to AAV9 in each cell line. (**F**) The binding kinetics between each capsid and full-length human TfR1 were assessed by BLI. AAVX probes were loaded with capsid and human TfR1 was used as an analyte. Sensorgram curve fits (black dash-dot lines) were generated by applying global 2:1 exponential association and decay models.

### BI-hTFR1 binds the TfR1 apical domain

To evaluate whether binding to TfR1 is necessary for the increased transduction of brain endothelial cells by BI-hTFR1, we assessed hCMEC/D3 transduction in the presence of two anti-TfR1 antibodies: OKT9, a monoclonal antibody that binds the TfR1 apical domain (*26*) or AF2474, a polyclonal antibody that competes with Transferrin (Tf) for TfR1 binding. We found that transduction was significantly inhibited by increasing concentrations of OKT9 but not by AF2474 (Fig. 2A), suggesting that an interaction with the human TfR1 apical domain was required for the enhanced transduction mediated by BI-hTFR1 in these cells. Consistent with this result, OKT9 directly competed with BI-hTFR1 for binding to purified human TfR1 (fig. S2B). Notably, binding to the TfR1 apical domain is a common feature of several antibody-based BBB shuttles that have been tested in humans and likely positions the binding site away from the Tf binding site. This is important because surface TfR1 is thought to be mostly occupied by Tf based on the typical 10–20 µM blood concentration of iron-bound holo-Tf (*27*) and its sub nanomolar binding affinity for TfR1 (fig. S2C). Given the double digit nanomolar affinity of BI-hTFR1 for human TfR1, Tf would be predicted to outcompete BI-hTFR1 for binding to TfR1 if their binding epitopes overlap. Using BLI, we found that BI-hTFR1 binds human TfR1 with similar kinetics whether in the presence or absence of a receptor-saturating concentration of holo-Tf (Fig. 2B and fig. S2D) and that BI-hTFR1 is capable of binding a preformed complex of human TfR1 and Tf (Fig. 2C). Consistent with the BLI results, OKT9, but not Tf, inhibited the association of BI-hTFR1 with hCMEC/D3 cells and its colocalization with human TfR1 (Fig. 2D and fig. S3).

**Fig. 2.**
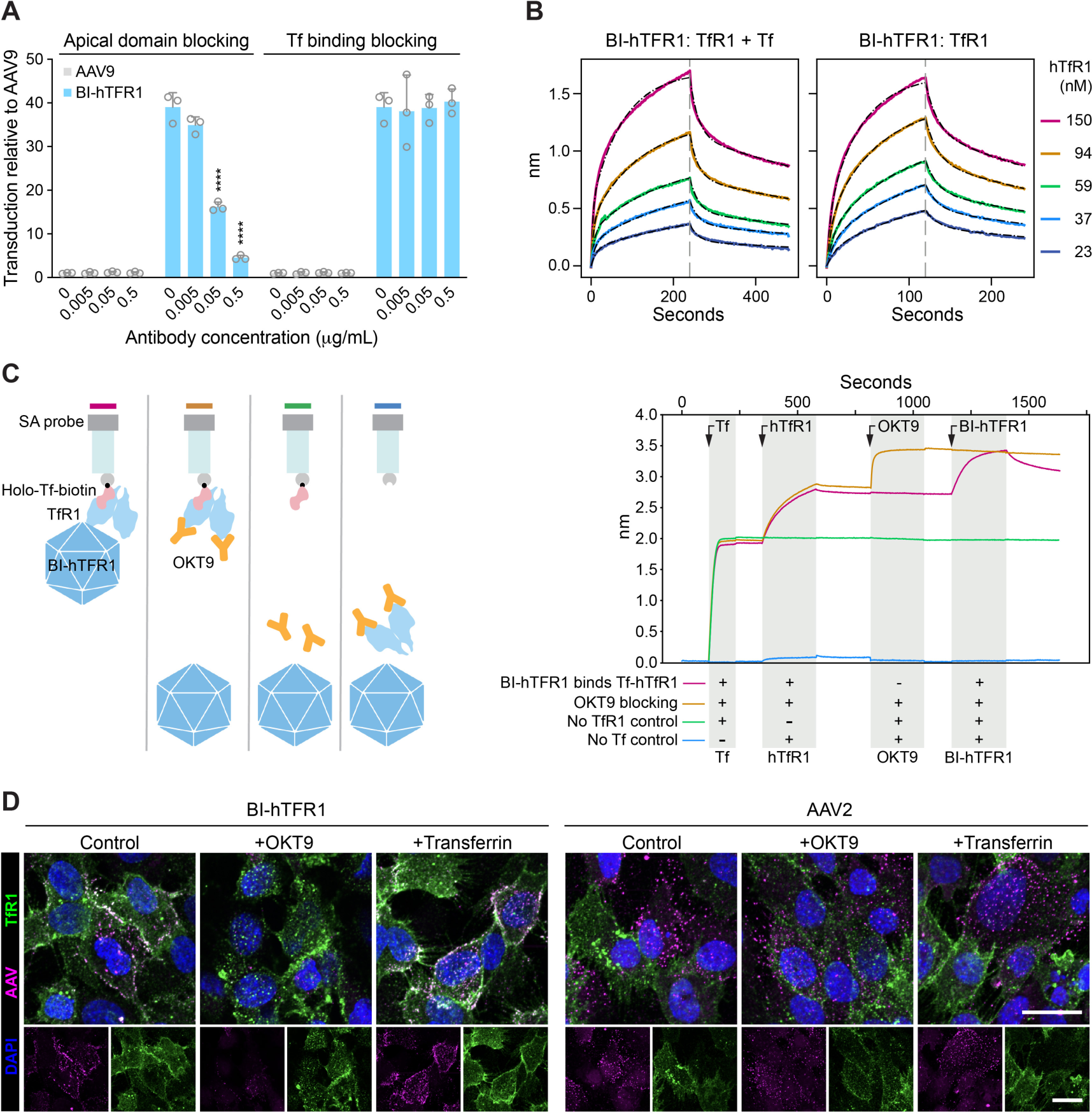
Human TfR1-targeted capsids transduce human brain endothelial cells via interactions with the apical domain of human TfR1. (**A**) The plots show transduction (Luciferase, normalized RLU) of hCMEC/D3 cells incubated with 3 x 10^8^ vg/mL of the indicated AAV and the specified concentrations of the OKT9 or AF2474 antibody (two-way ANOVA using the no antibody control for each condition as the main comparison group with Bonferroni multiple comparison correction: **** indicates *p* ≤ 0.0001; *n* = 3 replicates, error bars indicate ± SEM). (**B**) The effect of Tf on BI-hTFR1 binding to full-length human TfR1 was assessed by BLI. AAVX probes were loaded with AAV9 or BI-hTFR1. Human TfR1 that either had or had not been pre-incubated with 300 nM holo-Tf was used as an analyte. 2:1 binding model curve fits (black dash-dot lines) are shown. (**C**) Biotinylated holo-Tf was immobilized on streptavidin-coated BLI probes (SA probe), introduced first into human TfR1, and then into buffer with or without the OKT9 antibody, and finally into BI-hTFR1 virus particles. Segments shaded in gray highlight the indicated association step. (**D**) BI-hTFR1 or AAV2 was incubated with hCMEC/D3 cells at 50,000 vg/cell for one hour at 4°C, with or without OKT9 (1 µg/mL) or Tf (1 µg/mL), and immunostained for AAV and TfR1. Scale bars = 15 µm. AAV9 binding to hCMEC/D3 cells was rarely detected therefore AAV2 was used as a control (fig. S3).

### Active trafficking of BI-hTFR1 across a human BBB model

To test AAV transport across a human endothelial cell barrier, we established a BBB transwell model using the hCMEC/D3 cell line and compared the amount of BI-hTFR1 that was transported across the endothelial barrier relative to AAV9 and AAV2. To minimize the effect of well-to-well variation, we mixed BI-hTFR1, AAV9, and AAV2 carrying barcoded genomes that were individually identifiable by qPCR (table S1), added this mixture to the media of the upper chambers which were maintained at 37°C or 4°C, and measured transport to the lower chamber three hours later (Fig. 3A). At 4°C, ATP-dependent transcytosis is suppressed but passive transport, at least in part due to paracellular crossing, can occur, providing a measurement of leakiness that is a known issue with the use of hCMEC/D3 transwell assays (*28*). As previously reported, AAV9 crossed the transwell barrier more efficiently than AAV2 at 37°C, and the number of vector genomes detected in the bottom chamber was significantly reduced at 4°C for both AAVs as is consistent with ATP-dependent transcytosis (*29*–*31*). Significantly more BI-hTFR1 was actively transported to the bottom chamber than AAV9 or AAV2 (Fig. 3B). Passive transport of BI-hTFR1 and AAV2 were reduced relative to AAV9, likely due to increased association with the cells.

**Fig. 3.**
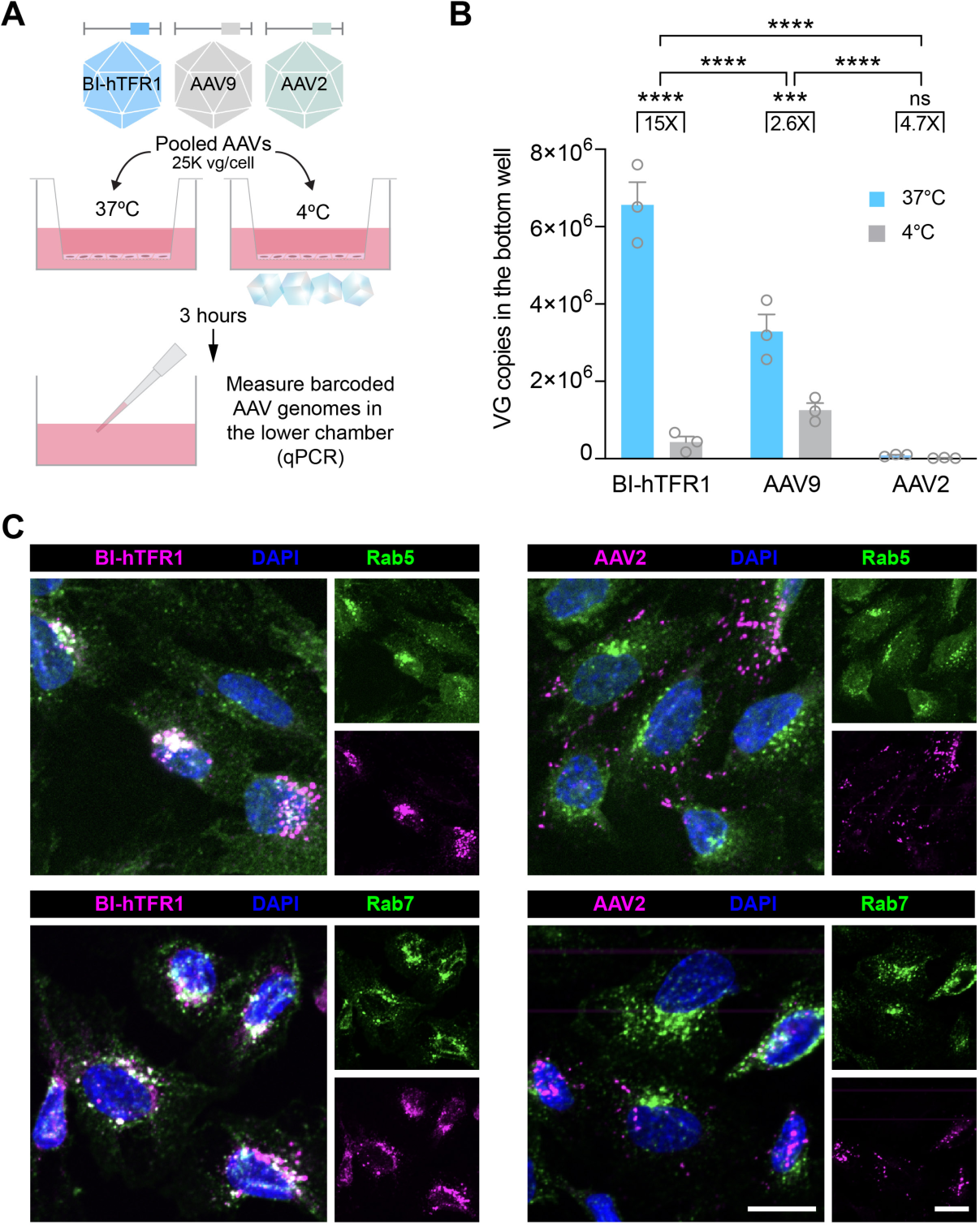
BI-hTFR1 is efficiently endocytosed and actively transported across human brain endothelial cells. (**A**) Schematic shows the pooled transwell BBB model experimental design. (**B**) The vector genomes in the bottom chamber were quantified by qPCR (two-way ANOVA with Bonferroni multiple comparison correction: **** and *** indicate *p* ≤ 0.0001 and ≤ 0.001 respectively; *n* = 3 transwell replicates, error bars indicate ± SEM). (**C**) BI-hTFR1 or AAV2 were incubated with hCMEC/D3 cells at 25,000 vg/cell for one hour at 37°C and stained for endosomal markers Rab5 and Rab7 as well as the AAVs. Scale bar = 15 µm.

RMT involves endocytosis, vesicular trafficking, and exocytosis (*32*). We explored whether BI-hTFR1 transcytoses hCMEC/D3 cells via the RMT pathway of TfR1 by assessing its colocalization with various organelle markers. We found that BI-hTFR1 partially colocalized with markers of the early and late endosomal pathway (Rab5 and Rab7) as well as the trans-Golgi network (TGN46), but did not extensively colocalize with markers of the cis-Golgi (Rcas1) or endoplasmic reticulum (KDEL) (Fig. 3C and fig. S4). In contrast, AAV2 was not highly colocalized with any of the endosomal or organelle markers with the exception of TGN46. Partial colocalization of both AAVs with TGN46 is consistent with the requirement for trafficking of AAVs to the trans-Golgi for transduction (*33*). Cell surface TfR1 clustering promotes clathrin-coated pit formation (*34*) and uptake into a subpopulation of endosomes enriched in Rab5 (*35*). This is consistent with our observation that BI-hTFR1 colocalized with Rab5 to a greater degree than AAV2. A smaller proportion of TfR1 colocalizes with Rab7 (*36*), which decorates endosomes associated with transcytosis (*37*) and the lysosomal degradation pathway. This is again consistent with our observation of a partial overlap between BI-hTFR1 and Rab7 localization in the cells. Our colocalization assays suggest that BI-hTFR1 engages TfR1 and is actively transported across the human BBB model via RMT. In contrast, AAV2 is predominantly trafficked to the trans-Golgi (*33*).

### CNS tropism of BI-hTFR1

Having established that BI-hTFR1 is more effective at transducing human brain endothelial cells via its interaction with TfR1 and is more efficiently transported across a human BBB model, we next sought to investigate whether BI-hTFR1 could cross the BBB *in vivo* by engaging human TfR1. To test this, we used *TFRC* knock-in (KI) C57BL/6J mice in which exons 4–19 of the mouse *Tfrc* encoding the extracellular domain were replaced by the corresponding region of human *TFRC* (Fig. 4A) (*38*). Levels of the mouse-human hybrid gene mRNA and protein products were found to be similar to those in WT mice (fig. S5). Therefore, we deemed these mice suitable for assessing the tropism of AAVs targeted to human TfR1. We intravenously injected BI-hTFR1 or AAV9:CAG-NLS-mScarlet-P2A-Luciferase-WPRE-SV40pA at 5 x 10^11^ vg/mouse into adult female C57BL/6J or *TFRC* KI mice (Fig. 4B). Three weeks post-injection, we observed enhanced biodistribution to and transduction of the brain and spinal cord by BI-hTFR1 in the *TFRC* KI but not in control C57BL/6J mice (Fig. 4C and D, Fig. 5A and B, and fig. S6). Increased biodistribution and transduction relative to AAV9 were not observed in any of the other organs that were assessed, indicating that the interaction with human TfR1 selectively enhanced the CNS tropism. Comparing the *TFRC* KI mice that received BI-hTFR1 to the C57BL/6J mice that received AAV9, we detected 6-fold and 12-fold more viral genomes; 54-fold and 43-fold higher viral mRNA transcript levels; 132-fold and 58-fold greater Luciferase enzymatic activity in the brain and spinal cord, respectively. As expected, AAV9 transduction of the CNS was not significantly affected by genotype.

**Fig. 4.**
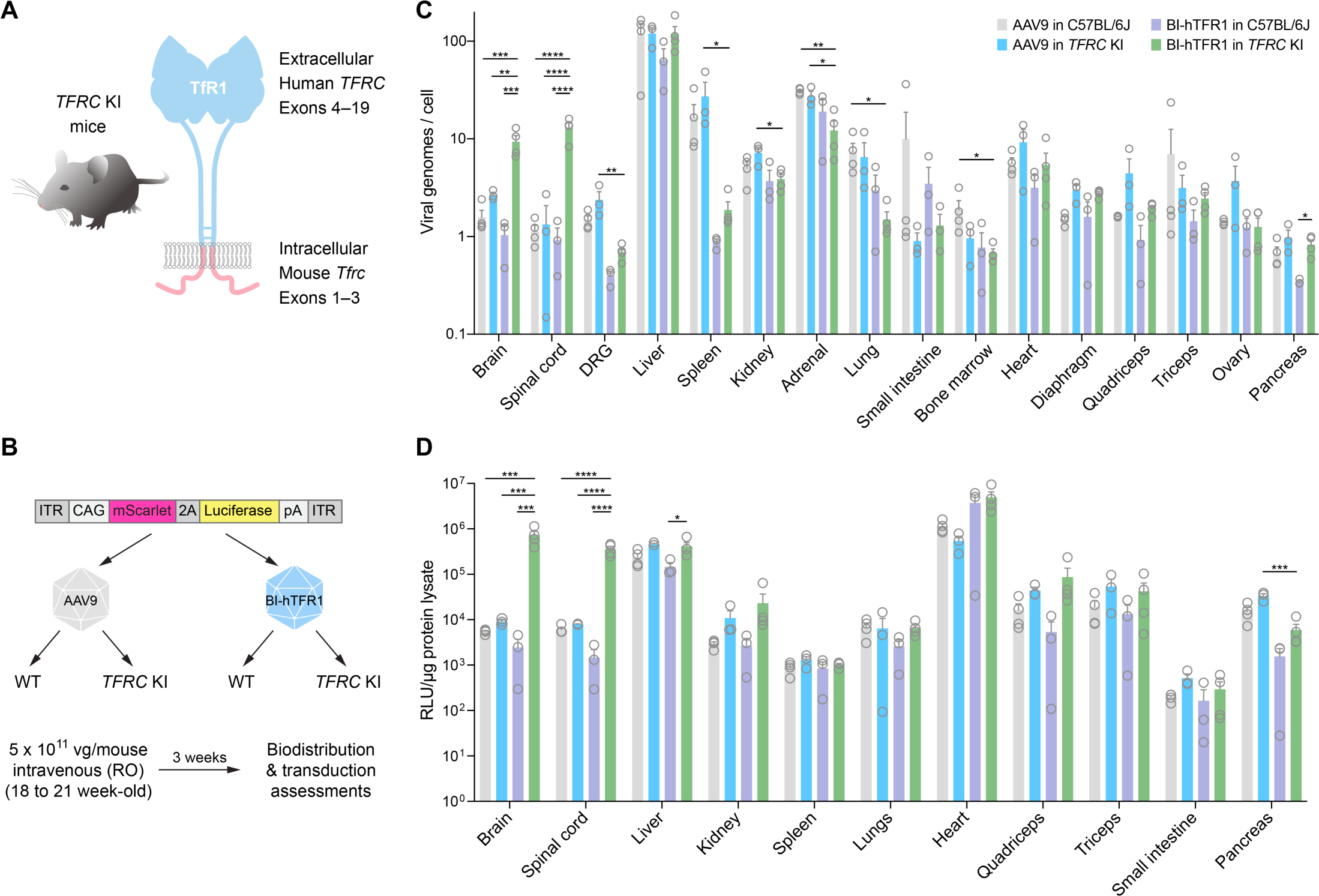
BI-hTFR1 efficiently delivers genes to the CNS of *TFRC* KI mice. **(A)** In *TFRC* KI mice, mouse *Tfrc* exons 4–19 encoding the extracellular region of TfR1 have been replaced by those of human *TFRC*. **(B)** BI-hTFR1 or AAV9 encoding CAG-NLS-mScarlet-P2A-Luciferase-WPRE-SV40pA were intravenously injected into adult female C57BL/6J or *TFRC* KI mice at 5 x 10^11^ vg/mouse. AAV9 in C57BL/6J and BI-hTFR1 in *TFRC* KI mice had *n* = 4 mice per group. AAV9 in *TFRC* KI and BI-hTFR1 in C57BL/6J had *n* = 3 mice per group. The (**C**) biodistribution reported as vector genomes per mouse genome and (**D**) Luciferase activity within different organs are shown at three weeks post-injection (two-way ANOVA using BI-hTFR1 in *TFRC* KI mice as the main comparison group with Bonferroni multiple comparison correction: ****, ***, **, and * indicate *p* ≤ 0.0001, ≤ 0.001, ≤ 0.01, and ≤ 0.05, respectively; each data point represents an individual mouse, error bars indicate ± SEM).

**Fig. 5.**
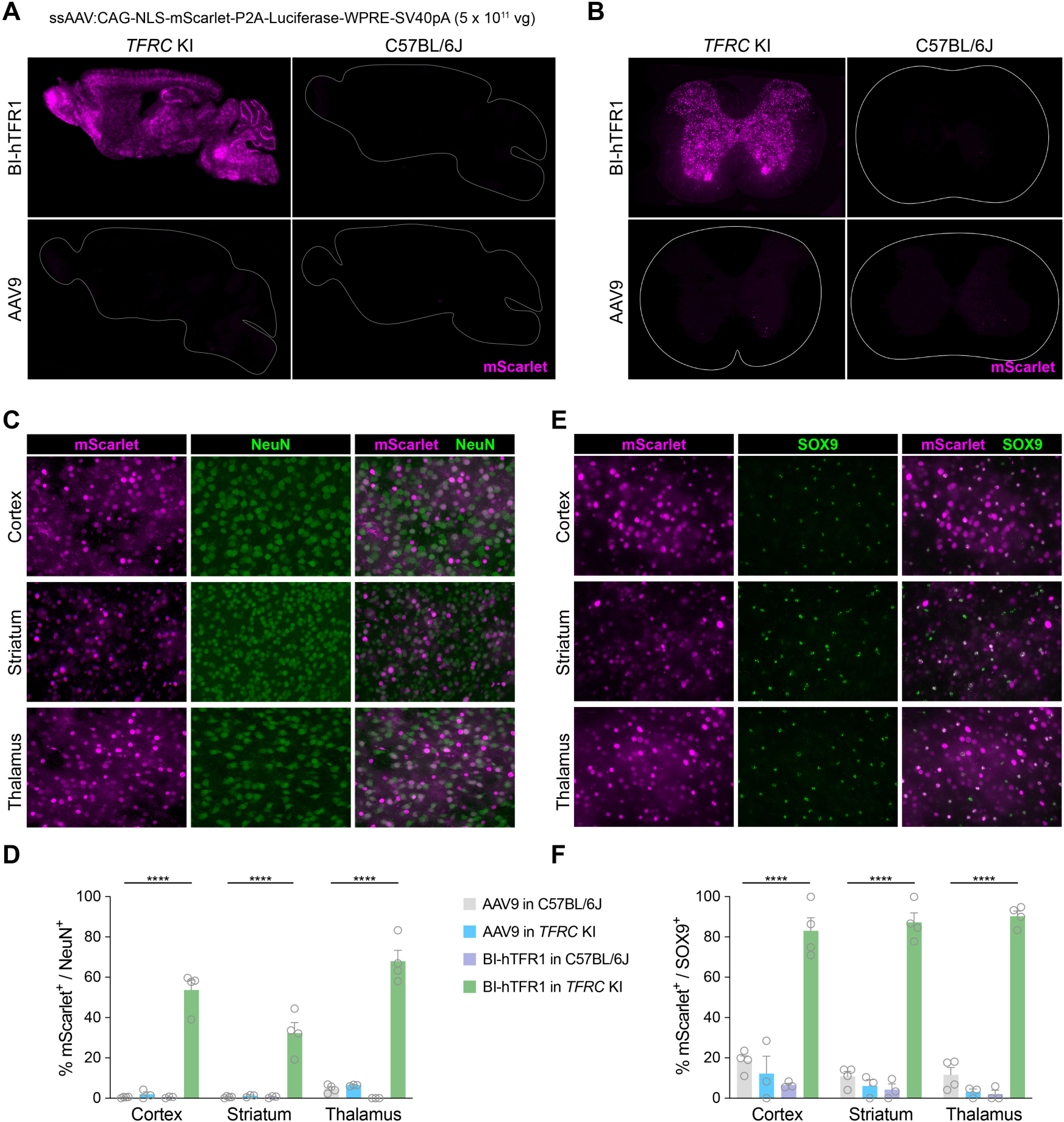
BI-hTFR1 efficiently transduces neurons and astrocytes throughout the CNS. Representative **(A)** whole brain and **(B)** spinal cord images from each group of mice at three weeks post-injection are shown. Representative images show cells transduced by BI-hTFR1 overlaid with **(C)** NeuN^+^ or **(E)** SOX9^+^ stained cells in the cortex, thalamus, and striatum of *TFRC* KI mice. The percentages of **(D)** NeuN^+^ neurons or **(F)** SOX9^+^ astrocytes that expressed mScarlet in the cortex, striatum, and thalamus are shown (two-way ANOVA using BI-hTFR1 in *TFRC* KI mice as the main comparison group with Bonferroni multiple comparison correction: **** indicates *p* ≤ 0.0001; each data point represents an individual mouse, error bars indicate ± SEM).

To evaluate the efficiency of transduction of specific cell types within the CNS, we assessed the fraction of cells positive for markers of neurons or astrocytes that expressed mScarlet. In *TFRC* KI mice, BI-hTFR1 transduced a mean of 54, 32, and 68% of NeuN^+^ neurons in the cortex, striatum, and thalamus, respectively (Fig. 5C and D). In contrast, in C57BL/6J mice, BI-hTFR1 transduced less than 1% of NeuN^+^ cells in the cortex and striatum and thalamus. AAV9 transduced 1–6% of NeuN^+^ cells in these same regions regardless of the mouse strain. BI-hTFR1 transduced more than 80% of the SOX9^+^ astrocytes in the cortex, striatum, and thalamus of *TFRC* KI mice, in comparison with 10% or fewer SOX9^+^ cells in C57BL/6J mice, or 20% or fewer SOX9^+^ cells transduced by AAV9 in C57BL/6J mice (Fig. 5E and F). Consistent with our biodistribution and transduction data, mScarlet levels in the livers of C57BL/6J and *TFRC* KI mice were comparable between AAV9 and BI-hTFR1 (fig. S6A and B). We observed a significant reduction of biodistribution to the DRGs in *TFRC* KI animals treated with BI-hTFR1 relative to AAV9 controls (Fig. 4C); DRG cell transduction by AAV9 or BI-hTFR1 in C57BL/6J or *TFRC* KI mice is shown in fig. S6C.

### BI-hTFR1 mediates GBA expression throughout the brain

We next evaluated BI-hTFR1’s ability to deliver the human gene Glucosylceramidase Beta 1 (*GBA1*) to *TFRC* KI mice. We built a single-stranded AAV genome containing a hybrid cytomegalovirus enhancer-chicken beta-Actin (CMV-CBA) promoter driving the expression of human *GBA1* with a C-terminal influenza virus hemagglutinin (HA) tag (Fig. 6A) (*39*). The genome was packaged into AAV9 or BI-hTFR1 and administered intravenously to *TFRC* KI mice at a dose of 1 x 10^14^ vg/kg, which is comparable to the weight-adjusted dose of the FDA-approved AAV9-based Zolgensma gene therapy for spinal muscular atrophy. Additionally, we administered BI-hTFR1:*GBA1* to a separate group of *TFRC* KI mice at a 20-fold lower dose (5 x 10^12^ vg/kg). For the untreated control mice, we used C57BL/6J mice, which possess the same genetic background as the *TFRC* KI mice. Three weeks post-administration, we detected ∼30 times more AAV genomes in the brains of mice treated with BI-hTFR1 as compared with those of mice that received AAV9 (Fig. 6B).

**Fig. 6.**
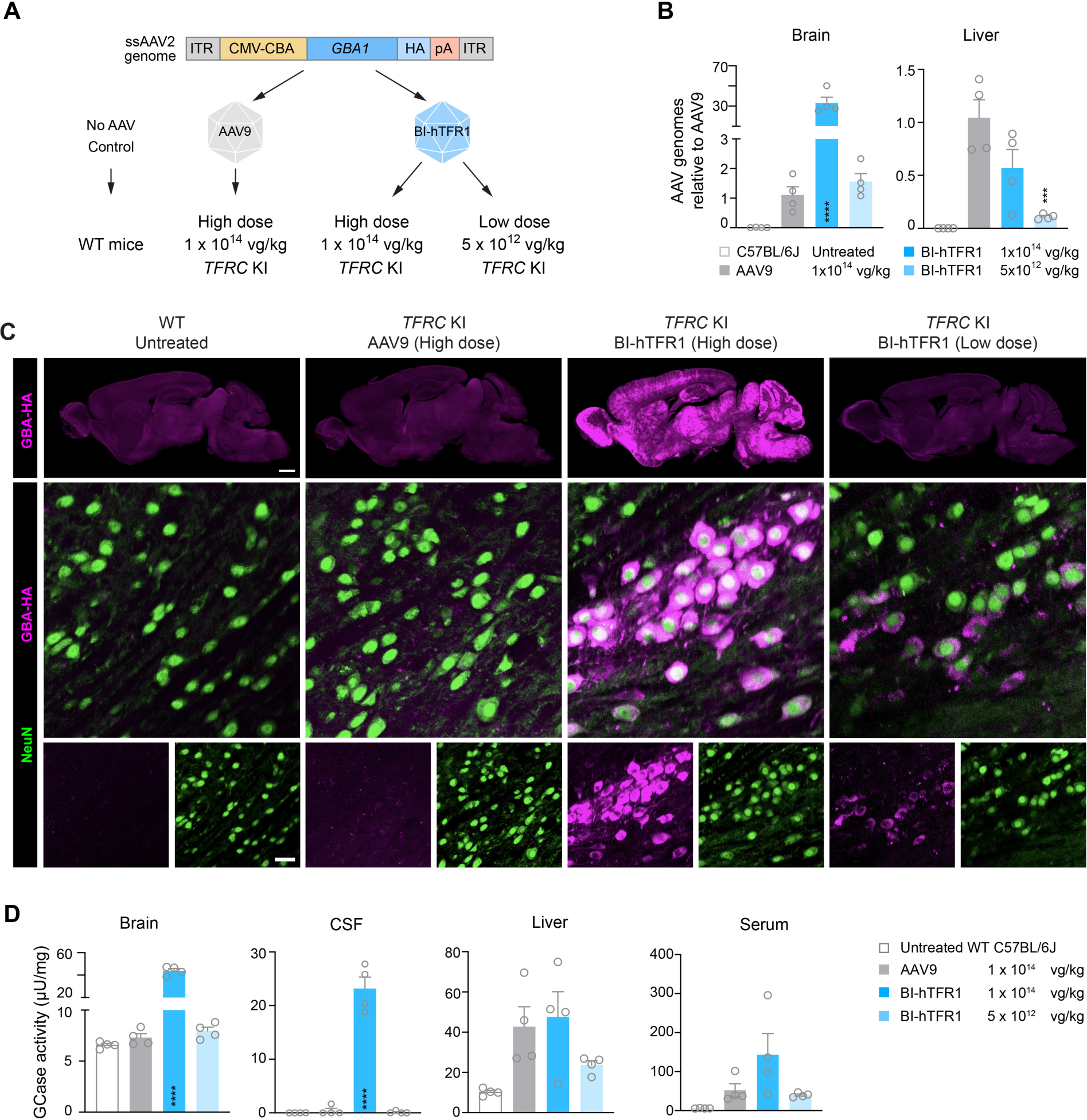
BI-hTFR1 efficiently delivered *GBA1* and increased GCase activity in the brains of *TFRC* KI mice. (**A**) Schematic of the experiment shows the ssDNA AAV genome expressing human glucocerebrosidase that was packaged into AAV9 or BI-hTFR1 and administered to *TFRC* KI transgenic mice at either 1 x 10^14^ vg/kg or 5 x 10^12^ vg/kg. (**B**) The biodistribution of AAV genomes found in brain and liver tissue relative to AAV9 is shown (one-way ANOVA using AAV9 as the main comparison group with Sidak’s multiple comparison post-hoc test: **** and *** indicate *p* ≤ 0.0001 and ≤ 0.001, respectively; each data point represents an individual mouse, error bars indicate ± SEM). (**C**) Sagittal brain sections (top) from the mice in each group show GBA-HA (magenta) in whole brain sagittal sections. Scale bar = 1 mm. Images of immunostaining (bottom) show neurons (NeuN, green) and GBA-HA (magenta) in the substantia nigra pars compacta. Scale bar = 25 µm. (**D**) GCase enzyme activity levels in brain and liver tissue homogenate, CSF, and serum are shown (one-way ANOVA using AAV9 as the main comparison group with Sidak’s multiple comparison post-hoc test: **** indicates *p* ≤ 0.0001; each data point represents an individual mouse, error bars indicate ± SEM).

We observed robust brain-wide expression of GBA-HA as assessed by HA immunostaining (Fig. 6C) and significantly elevated glucocerebrosidase (GCase) activity in brain tissue lysates and the CSF, compared to the mice injected with the high dose of AAV9 or control C57BL/6J mice (Fig. 6D, and fig. S7A and B). In the mice injected with the high dose of AAV9, we observed minimal GBA expression in the brain but similar amounts of viral genomes, GBA expression, and GCase activity in the liver compared to the mice injected with the high dose of BI-hTFR1 (Fig. 6B and 6D, and fig. S7C); as expected, in the mice injected with the low dose of BI-hTFR1, the amount of viral genomes in the liver was reduced relative to the high dose AAV9 group. Reductions in GCase activity in the brain and CSF have been reported to be associated with the progression of Parkinson’s disease (*40*, *41*), and previous studies suggest that *GBA1* delivery to the brain can treat neuronal α-synucleinopathy (*42–44*). Notably, in the mice that received the low dose of BI-hTFR1, we observed neuronal transduction in regions linked to motor symptoms of Parkinson’s disease such as the substantia nigra pars compacta, deep cerebellar nuclei, and other parts of the brainstem (Fig. 6C and fig. S7A) (*45*). Under the same imaging conditions, GBA-HA was not detected in the brains of the mice injected with the high dose of AAV9. This suggests that a 20-fold lower dose of BI-hTFR1 led to greater GBA levels in disease-relevant brain regions compared to AAV9.

## Discussion

In this study, we leveraged a well established mechanism—engagement of the apical domain of human TfR1— to shuttle an engineered AAV capsid, BI-hTFR1, across the BBB. Protein shuttles designed to bind human TfR1 to facilitate RMT are now in the clinic and are showing evidence of CNS entry (NCT04023994) and efficacy (*46*). The TfR1 apical domain is also a receptor target of natural viruses (*47–51*), indicating that this protein can be exploited for viral entry. Importantly, we demonstrated that BI-hTFR1 binds human TfR1 and pre-formed TfR1-Tf complexes with comparable kinetics and does not compete with holo-Tf for TfR1 binding. The ability to bind TfR1 in the presence of its natural ligand is critical because AAVs cannot be administered at doses capable of competing with Tf for TfR1 binding. The use of this validated transport mechanism bolsters our confidence that BI-hTFR1 can perform well as a CNS-targeting human gene therapy vehicle.

We engineered BI-hTFR1 to bind TfR1 by inserting a novel 7-mer TfR1 recognition sequence directly within surface loop VIII of the capsid. Because our approach did not require the introduction or conjugation of bulky protein domains as a means to retarget the AAV (*52–58*), we do not anticipate additional complexities in the manufacturing of BI-hTFR1 compared to AAV9, one of the top producing natural AAVs. Indeed, we consistently produced and purified BI-hTFR1 with final yields comparable to those of AAV9, and determined that BI-hTFR1 can be purified with affinity resins commonly used in AAV manufacturing (fig. S8)

As compared to natural AAV capsids such as AAV9 and AAV2 that are used in approved gene therapies, BI-hTFR1 demonstrated the remarkable ability to be actively transported across a human brain endothelial cell layer and to cross the BBB and transduce cells throughout the brains and spinal cords of *TFRC* KI mice after intravenous delivery. BI-hTFR1 exhibited a broad CNS tropism enhancement similar to those observed with engineered mouse BBB-crossing capsids, e.g., AAV-PHP.B, BI28, AAVF, and 9P31 that bind to mouse GPI-anchored proteins LY6A, LY6C1, or Car4 (*25*, *59*, *60*). By engaging highly abundant CNS endothelial cell surface proteins as new receptors, these capsids are able to efficiently cross the BBB and gain access to parenchymal cells.

A critical advantage of BI-hTFR1 and other capsids developed through target receptor binding screens is that their MOA is well established, thus their species tropism is predictable and can inform large animal experiments. This stands in contrast to capsids identified from conventional selections with unknown MOAs and therefore unknown potential to translate to other species including humans. We do not expect the human TfR1-targeting AAVs reported in this study to exhibit enhanced CNS tropisms in NHPs such as macaques and marmosets. Therefore, tests in these species would only be valuable in terms of profiling biodistribution, immune responses, and toxicity in the absence of the target, human TfR1. Knowing the precise target of the AAV capsid also aids the prediction of its function in human patients. For instance, there are no known missense variants with an allele frequency greater than 1 in a 1000 within the *TFRC* apical domain (*61*). Therefore, we predict that BI-hTFR1 binding to human TfR1 will not be impacted by common variations within the coding sequence of the human apical domain.

A second advantage of human receptor-targeting capsids is that gene therapies leveraging these vectors can be tested in KI mouse models possessing both the human target receptor and disease-relevant mutation(s). For instance, disease models are typically created using mice and, in the absence of a capsid with an enhanced CNS tropism that translates across species, studies will need to use a surrogate mouse BBB-crossing capsid to deliver and test their potentially therapeutic payloads in these disease models. With BI-hTFR1, the gene therapy product that would be delivered to human patients can be tested, without switching out the capsid for a surrogate, in a mouse disease model that has been crossed with *TFRC* KI mice. Using a therapeutically relevant payload, we found that BI-hTFR1 could deliver 30 times the number of AAV:*GBA1* genomes to the brains of *TFRC* KI mice and increase brain and CSF GCase activity as compared to AAV9. Even at a 20-fold lower dose, *GBA1* expression was detectable throughout the brain in BI-hTFR1:*GBA1* treated mice. However, the *TFRC* KI mice were WT with respect to *Gba1* and only the high dose of BI-hTFR1:*GBA1* led to increased brain and CSF GCase activity that were detectable above the endogenous mouse GCase activity.

Collectively, our results demonstrate that BI-hTFR1 can leverage the RMT activity of human TfR1 to cross the BBB and deliver genes widely and efficiently to neurons and glia throughout the CNS. These qualities provide strong support for the further exploration of BI-hTFR1 as a broadly applicable vehicle for human CNS gene therapies.

## Materials and Methods

### Animals

All procedures were performed as approved by the Broad Institute IACUC. C57BL/6J (000664) were obtained from The Jackson Laboratory (JAX). *TFRC* KI mice (C57BL/6-Tfr1tm1TFR1/Bcgen, B-hTFR1, 110861) were obtained from Biocytogen. Recombinant AAV vectors were administered intravenously via the retro-orbital sinus in young adult male or female mice. Mice were randomly assigned to groups based on predetermined sample sizes. Transgenic animals were genotyped using DNeasy Blood & Tissue Kit (Qiagen, 69504) using primers listed in table S1. Two mice were excluded from analysis in Fig. 4, Fig. 5, fig. S5, and fig. S6: The first was a mouse assigned to receive AAV9 in the *TFRC* KI group but was determined to be a WT by genotyping and western blotting. One mouse in the BI-hTFR1-treated C57BL/6J group displayed no evidence of transduction in the biodistribution, transduction, or Luciferase assays, and showed no fluorescent reporter signal in the liver or the brain. Experimenters were not blinded to the sample groups.

Mice were euthanized with Euthasol and transcardially perfused with PBS at room temperature followed by 4% paraformaldehyde (PFA) in ice cold PBS. Tissues were post-fixed overnight in 4% PFA in PBS and sectioned by vibratome. Immunohistochemistry (IHC) was performed on floating sections with antibodies diluted in PBS containing 5% donkey serum, 0.1% Triton X-100, and 0.05% sodium azide. Primary antibodies NeuN 1:500 (Invitrogen, MA5-33103), SOX9 1:250 (abcam, ab185966) were incubated at room temperature overnight. The sections were then washed and stained with secondary Alexa-conjugated antibodies Alexa Fluor 647 (Invitrogen, A-31573) and Alexa Fluor 555 (Invitrogen, A-21427) at 1:1000 for four hours or overnight.

### Plasmids

AAV9 and the BI-hTFR1 Rep-Cap plasmids were generated by gene synthesis (GenScript). The CAG-WPRE-hGH pA backbone was obtained from Viviana Gradinaru’s lab through Addgene (#99122). GFP, GFP-2A-luciferase, and mScarlet cDNAs were synthesized as gBlocks (IDT) or synthesized and cloned (GenScript).

Full-length *TFRC* cDNA expression and lentiviral plasmids were cloned by inserting the open reading frames of *TFRC* (human: NM_003234.4; mouse: NM_011638.4; marmoset: NM_001301847.1, macaque: NM_001257303.1) into the lentiviral backbone pLenti-EF-FH-TAZ-ires-blast (Addgene #52083) with EcoRI/SalI sites.

For the C-terminal Fc fusion protein, the coding sequence of the extracellular region of human TfR1 (residues 89–760) were amplified by PCR and inserted into pCMV6-XL4 FLAG-NGRN-Fc (Addgene #115773) with EcoRV and XbaI sites. The signal peptide H7 (ATGGAGTTTGGGCTGAGCTGGGTTTTCCTCGTTGCTCTTTTTAGAGGTGTCCAGTGT) was introduced to N terminal of the *TFRC* sequence to direct protein secretion.

For the N-terminal Fc fusion protein, the coding sequence of the extracellular region of human TfR1 (residues 89–760) was amplified by PCR. An N-terminal Fc region with an N-terminal H7 signal peptide and a C-terminal GS linker was ordered from IDT. The desired region of plasmid pCMV6-XL4 FLAG-NGRN-Fc (Addgene #115773) was amplified by PCR. These three DNA fragments were assembled with NEBuilder® HiFi DNA Assembly (NEB E2621).

### Cell lines and primary cultures

HEK293T/17 (CRL-11268), Pro5 (CRL-1781), and CHO-K1 (CCL-61) were obtained from ATCC. hBMVECs cells (H-6023) were obtained from Cell Biologics and cultured as directed by the manufacturer. hCMEC/D3 cells were obtained from Millipore-Sigma (SCC066).

### Reconstitution of full-length TfR1

Full-length human TfR1 was reconstituted into Peptidiscs (PEPTIDISC LAB, dissolved at 2 mg/mL in 20 mM TrisHCL, pH 8). Briefly, HEK293 cells expressing C-terminally tagged full-length human TfR1 were harvested by centrifugation at 300 × g. Cells were lysed by sonication and fractionated by centrifugation at 15,000 × g to remove nuclei and cell debris and at 100,000 × g to pellet the cell membrane. Membrane was resuspended in Buffer A (TrisHCl, pH 7.5, 50 mM NaCl, 10% glycerol) to 10 mg/mL, and then supplemented with 0.8% n-Dodecyl-β-D-Maltoside (DDM, Goldbio, no. DDM5) at 4°C with gentle agitation to solubilize the membranes. Anti-FLAG M2 magnetic beads (Sigma-Aldrich) were incubated with the solubilized membrane and washed with buffer A with 0.02% DDM. Beads were incubated for five minutes at room temperature in a solution of 1 mg/mL Peptidisc peptide and then thoroughly washed in buffer A without detergent. Peptidisc-TfR1 was eluted in TrisHCl, pH 7.5, 150 mM NaCl, 0.1 mg/mL 3XFLAG peptide and analyzed by SDS-PAGE.

### Bio-Layer Interferometry

A GatorPrime instrument was used for the characterization of the binding dynamics for all BLI studies and all analysis was conducted with Gator Bio software v 2.10.4.0713. Experiments were run in black 384 well plates (Greiner Bio-One, 781906). All reagents and buffer wells were prepared with Q buffer and probes from Gator Bio.

For AAV capsid binding to full-length human TfR1, AAVX probes (Gator Bio) were equilibrated in Q buffer and then loaded with either BI-hTFR1, BI-hTFR1-2, BI-hTFR1-3, BI-hTFR1-4, or AAV9 by uniform thresholding to an 8 nm shift. Seven serial dilutions of full-length human TfR1-FLAG were associated to each capsid for 120 seconds and then dissociated in Q buffer for 120 seconds. Only the five highest concentrations are shown on sensorgrams. Three trials of the assay were performed. A probe incubated with Q buffer instead of human TfR1 was included in each trial for background subtraction.

For BI-hTFR1 and AAV9 binding to pre-incubated Tf with full-length human TfR1, assays and analysis were conducted as above but with 300 nM holo-Tf (Millipore Sigma T413) pre-incubated with each dilution of TfR1 for 15 minutes.

For kinetic analysis of capsid binding to TfR1, the sensorgram data from each of three trials using seven different analyte concentrations per capsid displayed clear multi-phasic binding and dissociation characteristics that could not be explained by a monovalent interaction. Therefore, we used the global 2:1 exponential association and decay model. However, we are not implying that the binding between the TfR1-targeting capsids and TfR1 is bivalent. The kinetic constants shown in fig. S2 should be interpreted with caution due to the following factors: First and foremost, the high density of binding epitopes on the AAV surface may lead to local increases in the effective concentration of the receptor analyte, which results in a higher apparent affinity (*62*). Second, TfR1 itself is bivalent, which could contribute to the aforementioned avidity artifact and also result in the bridging of closely loaded adjacent AAVs. While probes were loaded to less than 60% of their total binding capacity, the uniformity of loading cannot be controlled and a fraction of the measurements may result from 2:1 binding of the receptor dimers to adjacent capsids (*63*). Third, transitional binding states due to conformational changes or sequential binding events may contribute to our observation of multi-phasic binding kinetics.

The software-computed rate constants from these replicate fittings were subject to QC thresholding that included R^2^ > 0.99, Gator-computed error ratios of <25% of their respective kinetic constants, and fitting residuals <10%. In the cases of both BI-hTFR1-3 and BI-hTFR1-4, one replicate was eliminated due to a k_on_ error ratio of greater than 25%. Trials with QC threshold-passing data were averaged and their standard error was taken.

For the tandem competition binding assays, a human TfR1 apical domain-targeting antibody, OKT9 (Thermo Fisher Scientific) was chosen as a competitor. The extracellular domain of human TfR1 (residues 89–760) was recombinantly expressed as an N-terminal Fc-fusion protein (Fc-hTfR1) in Expi293 suspension cells and purified by gravity flow with Pierce^TM^ Protein A Agarose (Thermo Fisher Scientific). HFCII probes were loaded with Fc-hTfR1 by thresholding to a 1 nm shift. Fc-TfR1-loaded probes were associated in tandem, first to BI-hTFR1 until the signal reached approximately 2 nm, and subsequently to the OKT9 antibody at 250, 100, 50, or 25 nM or in Q buffer alone. The latter was included to demonstrate BI-hTFR1 dissociation in the absence of competitor OKT9. In parallel, a Fc-hTfR1 loaded probe was associated first in Q buffer alone and next in 250 nM of the OKT9 antibody to demonstrate maximum antibody binding in the absence of capsid. The two association phases were 240 and 360 seconds, respectively.

For the triple tandem binding assays involving holo-Tf, holo-Tf was biotinylated for two hours on ice with an EZ-Link^TM^ Sulfo-NHS-LC-LC-Biotin kit (Thermo Fisher Scientific) and then buffer exchanged into PBS with a 0.5 mL Pierce Protein Concentrator PES column with a 10K MWCO to remove unreacted biotin. Streptavidin (SA) probes were equilibrated in Q buffer and then loaded to a 2 nm shift with biotinylated holo-Tf or were incubated in Q buffer to serve as a no-ligand control. Three of the probes were submitted to sequential binding steps for 240 seconds each. The probes were incubated in Q buffer for 240 or 120 seconds between each binding step to demonstrate <10% baseline drift (i.e., loss of binding) before binding the next molecule. In channel 1, the holo-Tf-loaded probe was associated first with 150 nM full-length human TfR1, next with Q buffer only, and finally with BI-hTFR1. In channel 2, the holo-Tf loaded probe was associated with TfR1, then 200 nM OKT9 antibody, and finally with BI-hTFR1 capsid. In channel 3, the holo-Tf-loaded probe was associated with Q buffer alone, then the OKT9 antibody, and finally with BI-hTFR1. In channel 4, the unloaded probe was associated with TfR1, then OKT9 antibody, and finally with BI-hTFR1 to reveal any nonspecific binding to the probe.

For full-length hTfR binding to human holo-Tf, MFC probes were equilibrated in Q buffer and then loaded with anti-Tf monoclonal antibody 12A6 (Thermo Fisher Scientific) to a 1 nm shift. Next, holo-Tf was loaded as ligand to saturation. Serial dilutions of full-length peptidisc-reconstituted hTfR1-FLAG were associated with holo-Tf for 120 seconds and then dissociated in Q buffer for 120 seconds. Three trials of the experiment were performed. A probe incubated with Q buffer alone was included in each trial for background subtraction in sensorgram curve fittings and kinetics analysis. For subsequent kinetics analysis, the sensorgram data from each of three trials were fitted with global 1:1 exponential association and decay models. The software-computed rate constants from these replicate fittings were subject to QC thresholding that included R^2^ > 0.99, computed error ratios of <10% of their respective kinetic constants, and fitting residuals <10%. Kinetics values were averaged and their standard error was taken.

### Lentivirus production

Lentivirus was produced with a third-generation lentivirus system by cotransfection of three packaging plasmids (pMDLg-RRE, pRSV-Rev, and pVSV-G) and vector plasmid (pLenti-EF-FH-TAZ-IRES-blast, a gift from Yutaka Hata, Addgene plasmid # 52083) encoding the entire coding sequences of *TFRC* or *Tfrc* (human: NM_001128148; macaque: NM_001257303; marmoset: NM_001301847; or mouse: NM_011638.4) at a ratio of 4:1:1:6 in HEK293T/17 cells. Lentiviruses were harvested from the media three days after transfection and filtered with 0.45 µm PES to remove cell debris.

### Exogenous TfR1 expression in CHO cell lines

CHO cell lines expressing human, macaque, or marmoset *TFRC* or mouse *Tfrc* were established via random lentiviral integration. Briefly, CHO cells were seeded at 150,000 cells/well in 12-well plates and transduced with lentivirus-containing media for 48 hours, then transferred to 10 cm dishes for selection with 10 µg/mL blasticidin for one week followed by maintenance in media with 1 µg/mL blasticidin. To validate expression of the desired construct, RNA was extracted from cells using the Qiagen RNeasy Kit. Maxima H Minus Reverse Transcriptase (ThermoFisher, EP0753) and oligo dT were used for cDNA synthesis. Species-specific primers were used for qPCR (table S1). CHO cells expressing TfR1 from each species were also assessed by immunostaining with two anti-TfR1 antibodies: ab84036 (Abcam) and OKT9 (eBioscienceTM) (fig. S1).

### AAV production and titering

The AAV9 variant capsids carrying CAG-NLS-mScarlet-P2A-Luciferase-WPRE-SV40pA or CMV/CBA-*GBA1*-HA-pA were either produced in suspension followed by iodixanol purification and titered as previously described (*64*) or produced using adherent cells as previously described (*65*). Production, purification methods, and post-purification yields are provided in figure S8. For BLI experiments, AAV vectors were produced in adherent HEK293T cells and purified by static binding (3 to 10 plates per preparation). At 72 hours post-transfection, the cells and supernatant (60–200 mL per prep) were collected in 125–500 mL shake flasks, incubated with 0.1% Triton X-100, 2 mM MgCl_2_ and 25 U/mL benzonase (Sigma, E1014-25KU) for 90 minutes at 37°C, and centrifuged for 10 minutes at 3724 ×g for clarification. The clarified lysates were incubated with 150 µL POROS AAV9 resin (Thermofisher, A27353), at 37°C for 90 minutes, under agitation (1.14 ×g). The mixes were loaded into detergent removal spin columns (Thermofisher, 87778) and supernatants were discarded using a vacuum manifold. POROS AAV9 beads were rinsed three times with 5 mL PBS, using a vacuum manifold. AAV vectors were eluted in 15 mL Falcon tubes, with 2 mL elution buffer (0.1 M Glycine, pH2.5) and neutralized with 500 µL 1 M Tris, 0.5 M NaCl, pH 8. AAVs were dialysed against 3 × 2 L of PBS (with calcium and magnesium) supplemented with 0.001% Pluronic F-68, using float-a-lyzer G2 dialysis cassettes with a MWCO of 8–10 Kd (Repligen, G235067), or buffer exchanged and concentrated in the same final formulation, using centrifugal filters with a MWCO of 100 Kd (Thermofisher, 88532).

### Affinity purification of BI-hTFR1

Affinity purification was performed using POROS CaptureSelect AAVX and AAV9 resins bought as pre-packed 1 mL columns (Thermo Fisher Scientific, A36652 and A36650). Prior to affinity purification, lysate was clarified using a Clarisolve CS40 depth filter (Millipore Sigma, CS40MS02H1) and an Opticap XL2 Polysep sterilizing 0.22 µm filter (Millipore Sigma, KGW3A02TT3). Clarified lysate was then frozen at -20°C until processing. Clarified lysate was warmed up at room temperature at least 12 hours prior to affinity purification and then sterile filtered using a vacuum bottle filter (VWR International LLC, 28199-778). AAVX and AAV9 columns were attached to an AKTA Pure 25 L chromatography system (Cytiva, 29018224) containing an auxiliary sample pump S9 (Cytiva, 29027745). The chromatography column was pre-equilibrated with five column volumes of deionized H_2_O and five column volumes of equilibration buffer (50 mM Tris, 150 mM NaCl, 0.01% Pluronic F-68, pH 7.5) prior to application of the AAV-containing clarified lysate at a flow rate of 0.5 mL/min. 195–240 column volumes of clarified lysate were loaded onto the columns at a flow rate of 0.5 mL/min. After sample application, the column was washed with five column volumes of equilibration buffer and high salt wash buffer (50 mM Tris, 500 mM NaCl, 0.01% Pluronic F-68, pH 7.5). The bound AAV was eluted using five column volumes of buffer A (100 mM Citric Acid, 75 mM NaCl, 0.5 M L-Arginine, 0.01% Pluronic F-68, pH 3.0) for POROS CaptureSelect AAV9 or five column volumes of Buffer B (100 mM Citric Acid, 75 mM NaCl, 0.01% Pluronic F-68, pH 2.5) for POROS CaptureSelect AAVX. Isocratic elution was performed in upflow at a flow rate of 0.5 mL/min and samples were collected based on peak fractionation. Resin was then washed with five column volumes of equilibration buffer. Following re-equilibration, strip phases were performed using five column volumes of 0.1 M phosphoric acid and high pH strip buffer (20 mM Tris, 0.5 M L-Arginine, 0.2 M NaOH, 0.5M NaCl, pH 10.8). Strip phases were collected using the fraction collector. Resin was then washed with five column volumes of equilibration buffer, deionized H_2_O and 20% ethanol. Neutralized samples were stored at -20°C until titration.

### Binding assays

CHO cells stably expressing human, macaque, or marmoset *TFRC* or mouse *Tfrc* were seeded at 30,000 cells per well; hCMEC/D3 cells were seeded at 30,000 cells per well; and hBMVEC cells were seeded at 15,000 cells per well. At 24 hours post-seeding, each well was subjected to a media change with the fresh cold media that contained each AAV variant carrying CAG-NLS-mScarlet-P2A-Luciferase-WPRE-SV40pA at 3,000, 6,000, or 12,000 vg/cell for CHO, hCMEC/D3, or hBMVEC cells, respectively. The plate was then maintained at 4°C with gentle rocking for one hour, followed by five phosphate buffered saline (PBS) washes. Cells were treated with Proteinase K at 56°C for one hour, followed by heat inactivation at 95°C for 10 minutes. Total DNA extraction was performed using the DNeasy kit (Qiagen). The DNA was diluted 1:20 and subjected to qPCR with the mScarlet-qPCR-F and mScarlet-qPCR-R primers (table S1).

### Transduction assays

CHO cells stably expressing human *TFRC* or macaque, marmoset, or mouse *Tfrc* were seeded at 100,000 cells per well; hCMEC/D3 cells were seeded at 10,000 cells per well; and hBMVEC cells were seeded at 5,000 cells per well. At 24 hours post-seeding, each well was subjected to a media change with the fresh media that contained the AAV variant carrying CAG-NLS-mScarlet-P2A-Luciferase-WPRE-SV40pA at 3,000, 6,000, or 12,000 vg/cell for CHO, hCMEC/D3, or hBMVEC cells, respectively, and the plate was maintained at 37°C with 5% CO_2_ for 24 hours. Transduction was measured using the Britelite Plus reporter gene assay system (PerkinElmer, 6066766). Luciferase activity was reported in relative light units (RLU) as raw data or normalized to AAV9 controls.

### Antibody inhibition assays

The hCMEC/D3 cells were seeded at 7,500 cells per well. Two days later, 100 µL of media with 3 x 10^8^ vg/mL AAV and the specified concentration of OKT9 or R&D AF2474 antibody was transferred to each well. The plate was maintained at 37°C with 5% CO_2_ for 24 hours. Transduction was measured using the Britelite Plus reporter gene assay system.

### Transwell assays

hCMEC/D3 cells (under nine passages) were maintained in EGM2-MV media with VEGF (Lonza, CC4147). Falcon® Permeable Supports for 24-well Plates with 1.0 µm Transparent PET Membranes (Corning, 353104) were prepared by coating with collagen type I (Millipore, 08-115) diluted 1:50 in PBS and incubating at 37°C for two hours. On Day 0, cells were plated at 30,000/cm^2^ on the membrane and grown for two days in 200 µL of EGM2-MV media with VEGF on the top of the membrane and 600 µL of media in the lower chamber. On Day 2 and Day 4, the media at both the top and bottom of the transwell was changed to EGM2-MV media without VEGF, and the media at the bottom of the transwell was changed to Astrocyte Conditioned Media (Sciencell, 1811). Starting from Day 4, TEER values were measured using the EVOM2 meter (World Precision Instruments). TEER values were calculated as the ohmmeter readout in wells with cells minus the ohmmeter readout in wells without cells, multiplied by the surface area in cm^2^ units.

Transwell permeabilities using this process were previously validated by measuring the permeability of Dextran FITC 4k (Sigma, 46944), 40k (Biotium, 76221-470), and 70k (Biotium, 76221-460). TEER values peaked on Day 6, reaching values >10 Ω/cm^2^, which is when AAVs were added to the transwells. BI-hTFR1, AAV2, and AAV9 with capsid-specific barcoded transgenes were pooled and added to the top well of the transwell at 25,000 vg/cell (each AAV). After three hours of incubation at 37°C or 4°C, 22 µL of media was extracted from the bottom chamber. To detect the amount of AAV in the media, extracts were treated with DNaseI (NEB, M0303S) for 15 minutes at room temperature, then diluted in TE buffer (5 mM EDTA), and denatured at 70°C for 10 minutes to deactivate DNaseI. The extracts were then diluted in 5% Tween 20 with UltraPure™ Salmon Sperm DNA Solution (ThermoFisher, 15632011). The amount of AAV that passed through the transwell was measured by qPCR using a standard curve of known AAV quantities derived from AAV-containing media applied to the top well.

### Immunofluorescence staining and colocalization in hCMEC/D3 cells

For the imaging of BI-hTFR1 or AAV2 binding to hCMEC/D3 cells with or without OKT9 or Tf-647, BI-hTFR1 and AAV2 were added to hCMEC/D3 cells in optical plates (PhenoPlate 96-well, PerkinElmer, 6055302) at 50,000 vg/cell, while Tf-647 (ThermoFisher, T23366) and purified TfR1 antibody (OKT-9, AAT Bioquest, 10713000) at a 1:200 dilution were added to wells to reach a final concentration of 1 µg/mL each. Cells were incubated for one hour at 4°C, rinsed three times with PBS, fixed with 4% paraformaldehyde, and blocked (Avidin/Biotin Blocking Kit, SP-2001, Vector Laboratories) prior to applying Anti-AAV9 (1:200 dilution, ThermoFisher, 7103332100) to the wells treated with BI-hTFR1 or the wells with no virus, or Anti-AAVX (1:200 dilution, ThermoFisher, 7103522100) to the wells treated with AAV2. After three PBS washes to clear away unbound primary antibodies, Streptavidin 555 (1:500 dilution, Thermofisher, S21381) and Alexa Fluor© 488 (1:500 dilution, JacksonImmunoResearch, 115-545-003) were applied for the visualization of AAVs and TfR1, respectively.

To image AAV colocalization with Tf-647, BI-hTFR1 or AAV2 were added to hCMEC/D3 cells in optical plates at 25,000 vg/cell with Tf-647 (1 µg/mL). Cells were incubated for one hour at 4°C, followed by immunostaining as described above.

To image AAV and TfR1 colocalization, BI-hTFR1 or AAV2 were added to hCMEC/D3 cells in optical plates at 25,000 vg/cell. Cells were incubated for one hour at 37°C, prepared for immunostaining as described above but with the addition of a permeabilization step prior to the blocking of endogenous biotin to visualize TfR1. Streptavidin 555 (1:500 dilution, Thermofisher, S21381) and goat anti-mouse 647 nm (1:500 dilution, 115-605-062, Jackson ImmunoResearch) were applied.

For organelle colocalization, the following conjugated reagents were applied: Rab5 ((D-11) FITC, 1:100 dilution, Santa Cruz Biotechnology, sc-46692), Rab7 (1:400 dilution, abcam, ab198337), KDEL (1:500 dilution, abcam, ab184819), RCAS1 (1:200 dilution, Cell Signaling Technology, 12290), and TGN46 (1:50 dilution, abcam, ab50595). For unconjugated antibodies (RCAS1 and TGN46), goat anti-rabbit 488 (Alexa Fluor® 488 AffiniPure Goat Anti-Rabbit IgG (H+L), 1:500 dilution, JacksonImmuno, 111-545-144) was applied to visualize the staining. For wells requiring dual mouse antibodies (those co-stained with Rab5 and OKT9), cells were blocked for one hour with mouse-on-mouse blocking reagent (ReadyProbes™ Mouse-on-Mouse IgG Blocking Reagent, 1:30 dilution, ThermoFisher, R37621) between antibody staining procedures.

Images for all experiments were obtained with a 60X oil objective on a Nikon Ti-e inverted spinning disc confocal microscope, with the exception of TGN46 staining, which was performed using a 63X oil objective on a Zeiss LSM 900 confocal microscope. Within each experiment, images were taken with predetermined, optimized, and fixed exposure times to allow image comparisons.

### Imaging and tissue cell counting

Images of native mScarlet in the whole brain, cortex, striatum, thalamus, spinal cord, liver, or the dorsal root ganglion were taken on a Keyence BZ-X810. For images of whole brain sections, single plane tiled images were taken with autofocus mode on and stitched using the Keyence Analysis Software. For comparisons of native mScarlet fluorescence across groups, imaging settings were optimized for BI-hTFR1 in *TFRC* KI mice and then applied across all samples and all groups within each organ. Adjustments to the levels and gamma in the whole brain, cortex, striatum, thalamus, spinal cord, liver, and dorsal root ganglion images were performed using identical settings across all samples within each brain region or tissue type.

Stained sections were imaged on a Keyence BZ-X810 using sectioning mode with a Z-stack depth of 10 µm during acquisition. For cell counting, image settings were exposed to detect low mScarlet expressing cells in each sample group within each brain region. The Z-stack was reconstructed into a full-single-stack image using Keyence Analysis Software and exported as separate TIFs for each channel imaged. Images were then imported as input data into ilastik version 1.4.0 (*66*) where the Pixel Classification within the segmentation workflow was used. All color/intensity, edge, and texture features were selected and used during training. Training was performed to distinguish foreground and background pixels across images that represented the cortex, striatum, or thalamus in C57BL/6J or *TFRC* KI mice that had received BI-hTFR1 or AAV9 in NeuN or SOX9 stained brain sections. Predictions were then exported as probabilities in TIF format and then imported into CellProfiler 4.2.4 (*67*). A custom workflow was created to determine percent overlap between positively transduced cells and antibody markers. First, channels were aligned using the Align module. Second, the IdentifyPrimaryObjects module with Otsu two classes thresholding method was used to detect and segment foreground objects (positively transduced cells, NeuN-positive, or SOX9-positive cells) from background in the images. Third, foreground objects were then expanded by 2–5 pixels using the ExpandOrShrinkObjects module. Fourth, the MaskObjects module was used to determine the number of colocalized objects that overlap with a threshold of 40% overlap or more. A separate CellProfiler workflow was created where the IdentifyPrimaryObjects module was used to detect foreground and background objects (two class) and then the OverlayOutlines module was used to overlay an outline of detected foreground objects onto the raw input image of the mScarlet, NeuN, or SOX9 channel. The images were exported with the SaveImages module. The exported image was then used to verify if further training with the Pixel Classification in ilastik was needed in order to detect missed or improperly classified cells. An independent evaluator without prior knowledge of expected outcomes conducted a manual count of randomly selected images using the Photoshop Count Tool to determine the percent overlap of mScarlet-positive cells with NeuN or SOX9-positive cells in stained tissue. The results were evaluated to be within ± 5% deviation.

### Western blot

Purified N-terminal flag tagged full-length human and mouse TfR1 proteins or whole mouse brain lysates were separated on Bolt 4–12% Bis-Tris Plus gels and transferred onto nitrocellulose membranes. After incubation with anti-TfR1 antibodies (0.5 µg/mL ThermoFisher, Catalog # 13-6890; 0.25 µg/mL R&D, AF2474) or anti-ß-actin (1:5000; Abcam, AB20272) followed by incubation with a horseradish peroxidase (HRP)-conjugated secondary antibody. The detection of the HRP signal was by SuperSignal West Femto Maximum Sensitivity Substrate using a Bio-Rad ChemiDoc TM MP system #1708280.

### *In vivo* biodistribution, transduction, and Luciferase assays

BI-hTFR1 or AAV9 encoding CAG-NLS-mScarlet-P2A-Luciferase-WPRE-SV40pA were administered intravenously to 18 to 21-week-old female C57BL/6J or TFRC KI mice at a dose of 5 x 10^11^ vg/mouse. After three weeks, the mice were perfused with PBS. Tissue samples from the brain, liver, spinal cord, and DRGs were dropped fixed into 4% PFA. Remaining tissues, including additional samples of the brain, liver, spinal cord, and DRGs were collected snap frozen on dry ice and stored at −80°C. Biodistribution analysis: Samples were processed using a DNeasy 96 Blood & Tissue Kit (Qiagen, 69581). qPCR was performed for the mScarlet transgene and the mouse glucagon gene as previously described (see primer sequences in table S1) (*5*). For mRNA analysis, RNA was isolated using TRIzol (Invitrogen, 15596026) and purified using an RNeasy 96 Kit (Qiagen, 74171). cDNA synthesis was performed using Maxima H Minus Reverse Transcriptase (Thermo Scientific, EP0753). qPCR for mScarlet and mouse GAPDH was performed (primer sequences in table S1).

Luciferase assays: Protein was extracted from tissue samples using the lysis buffer T-Per with 1X Halt protease inhibitor (Invitrogen, 78430) while kept ice cold at all times. 10 µg of total protein was used to assess the reporter gene using the Britelite Plus reporter gene assay system.

### GBA1 *in vivo* delivery experiment

BI-hTFR1 or AAV9 encoding CMV/CBA-GBA1-HA-pA was intravenously injected into 28 to 30-week-old female TFRC KI mice at a dose of 1 x 10^14^ or 5 x 10^12^ vg/kg. After three weeks, cerebrospinal fluid (CSF) was collected from the cisterna magna of anesthetized mice as previously described (*68*). Briefly, the cisterna magna was surgically exposed and punctured with a sharpened glass capillary, and CSF was allowed to flow into the capillary. Approximately 6–10 µL of CSF was collected from each animal. Then, the thoracic cavity was exposed and 200–400 µL of blood was collected into a microtainer tube with serum separator additive (BD, 365967). The mice were then transcardially perfused with ice cold PBS. Brain, liver, and spinal cord tissue sections were drop fixed into 4% PFA. Additional tissue samples from these organs were collected and snap frozen at −80°C. After a 1.5-hour incubation at room temperature, the blood was centrifuged at 10,000 ×g for 90 seconds, and serum was collected and stored at −80°C. IHC was performed on fixed tissues as described in the Animals section of the Methods. GBA-HA was detected with an anti-HA primary antibody (Sigma, 11867423001) at a 1:50 dilution, followed by an AlexaFluor568-conjugated secondary antibody at a 1:500 dilution (Invitrogen, A78946). Vector genomes were quantified using qPCR primers targeting CMV and mouse GAPDH (table S1).

To measure GCase activity, whole tissue lysates were prepared. Tissue was thawed on ice, weighed, and homogenized in a 2× volume of ultra-pure distilled water with cOmplete protease inhibitor (Sigma, 4693132001) using the GenoGrinder (SPEX Sample prep). Homogenate was then centrifuged at 10,000 ×g for 2 minutes, transferred to a new, pre-chilled 1.5 mL Eppendorf tube, and incubated on ice for 15–30 minutes. Next, the samples were centrifuged at 12,000 ×g for 20 minutes. Supernatant was collected and stored at −80°C or used in subsequent analysis. A BCA assay (Thermo Fisher Scientific, 23225) was used to determine total protein concentration in the resulting lysates, as described by the product manufacturer. The SensoLyte Red Glucocerebrosidase (GBA) Activity Assay Kit (AnaSpec, AS-72259) was used to determine enzyme activity in serum and whole tissue lysate samples. Briefly, 20 µL of serum or the volume of tissue lysate containing 30 µg of protein was added into each well. Assay buffer was added to a total volume of 50 µL. Then, 50 µL of 1X enzyme substrate was added to each well. Samples were gently mixed and incubated for one hour at room temperature. The fluorescence signal at an excitation/emission of 570/610 nm was measured in end-point mode using the EnVision 2104 plate reader (PerkinElmer). A resorufin reference fluorescence standard curve was used to determine the amount of reaction product in each well from the measured relative fluorescence units (RFU). The resulting values were used to calculate enzyme activity, expressed in µU per mL of blood or µU per mg of protein. One unit of GCase activity is the amount of enzyme that generates 1 µmol of product per minute under the specified reaction conditions.

## Acknowledgments

We thank the members of the Deverman laboratory for discussions of the project. We also thank Nadine Elowe and Tonia Aristotelous for helpful advice with the BLI studies.

## Author contributions

Conceptualization: B.E.D., Q.H., K.Y.C.

Methodology: B.E.D., Q.H., K.Y.C., C.K., A.M., S.L., J.W.

Investigation: Q.H., K.Y.C., J.J., A.J.B., Q.Z., A.T.C., T.B., C.L., A.M., C.K., S.P., P.P.B., J.W.H., S.L., G.C., J.W., N.R.B.R., F.E.E., J.K.H., I.G.T., M.P.

Formal analysis: Q.H., K.Y.C., A.T.C., A.J.B., S.L., J.J., G.C., C.L., N.R.B.R., J.W., B.E.D.

Visualization: Q.H., J.J., S.L., G.C., K.Y.C., C.L., A.M., C.K. Y.A.C., J.W., N.R.B.R., B.E.D.

Funding acquisition: B.E.D.

Supervision: B.E.D., with support from Q.H., K.Y.C., Y.A.C.

Writing – original draft: Q.H., Y.A.C., B.E.D.

Writing – review & editing: all authors.

## Data and materials availability

All data are available in the main text or the supplementary materials. BI-hTFR Rep-Cap sequences and plasmids will be made available on Addgene at the time of publication. For pre-publication reagent requests please contact B.E.D.

## Supplementary Materials

**Figure S1.**
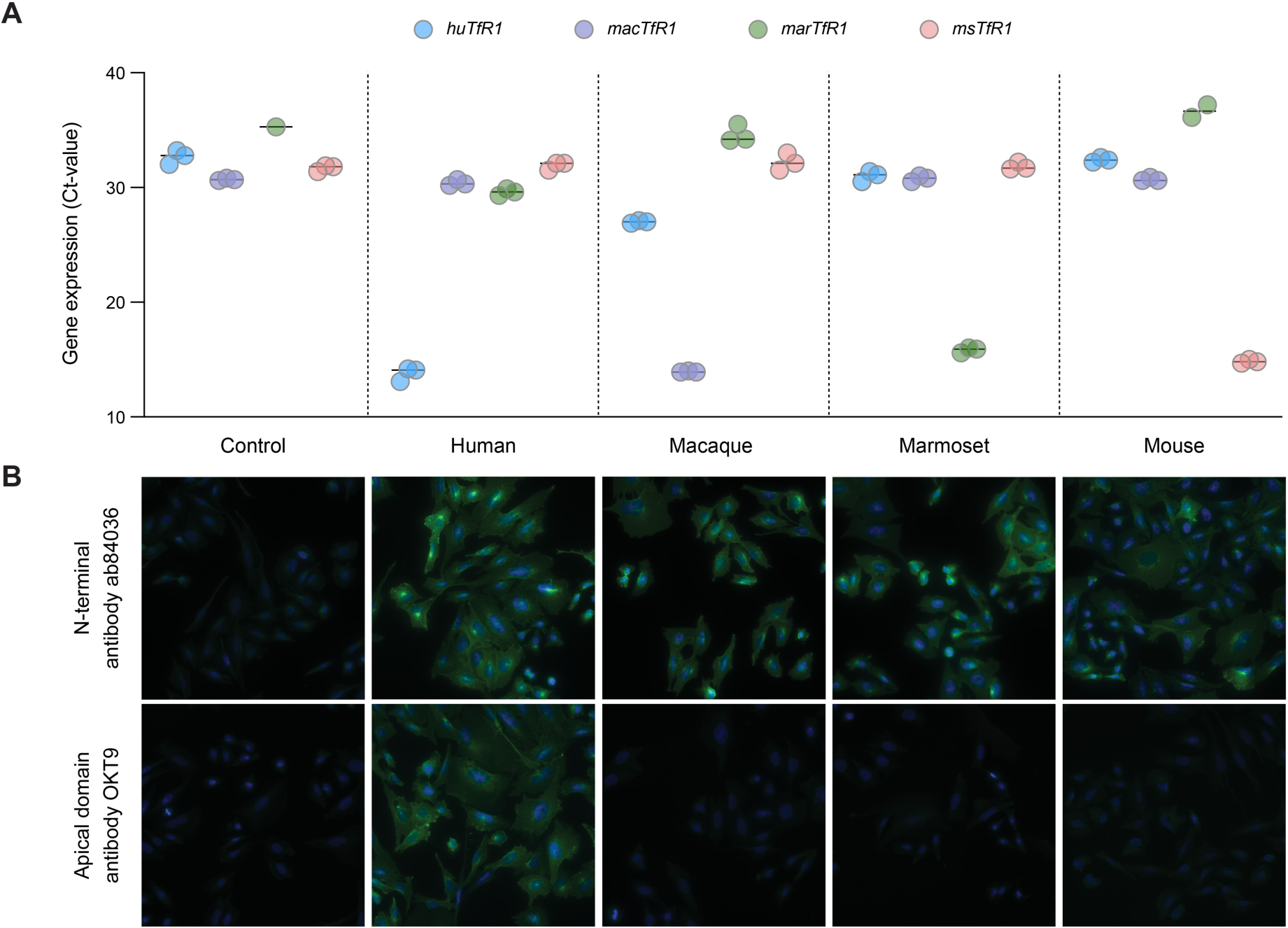
Validation assays confirm expression of TfR1 from the indicated species in stable CHO cell lines. (**A**) Species-specific primers (table S1) were used to assess *TFRC* or *Tfrc* mRNA expression levels by RT-qPCR. Each species primer set is shown with the indicated color (legend, top). The graph shows qPCR cycle threshold (Ct) values showing selective early cycle number amplification of the target species sequence. The bar indicates the mean Ct value (*n* = 3 replicates per cell line). (**B**) Images show immunofluorescence for TfR1. Each cell line was fixed, permeabilized, and stained with the indicated anti-TfR1 antibodies to assess the expression of exogenous TfR1.

**Figure S2.**
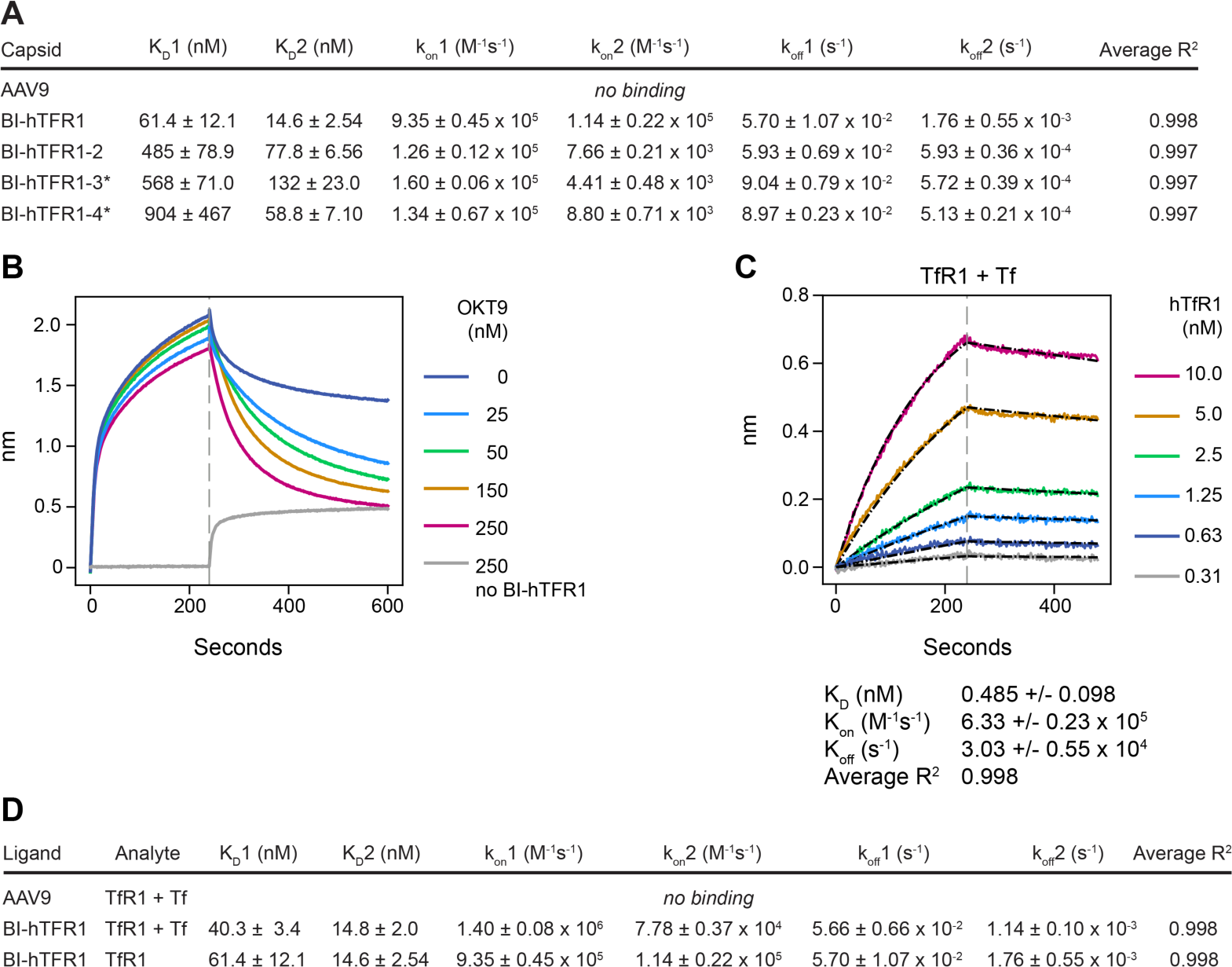
The binding between BI-hTFR1 and human TfR1 is inhibited by the apical domain binding antibody OKT9 but not by holo-Tf. (**A**) To assess binding between each AAV capsid and full-length human TfR1, AAVX probes were loaded with capsid and TfR1 was used as an analyte. Average kinetic constants and standard errors were computed from triplicate trials; * indicates a capsid for which values were computed from only two trials. (**B**) The sensorgram shows the association of BI-hTFR1 to the Fc-hTfR1-loaded HFCII probe and dissociation in the presence of the indicated concentration of OKT9. The gray curve shows OKT9 binding to Fc-hTfR1 in the absence of BI-hTFR1. (**C**) The binding kinetics between full-length human TfR1 and human holo-Tf were assessed by BLI. Mouse Fc (MFC) probes were loaded with an anti-transferrin antibody, 12A6. Holo-Tf was loaded onto the probe and associated with serial dilutions of TfR1 analyte. (**D**) The effect of Tf on BI-hTFR1 binding to full-length human TfR1 was assessed by BLI. AAVX probes were loaded with AAV9 or BI-hTFR1. Human TfR1 that either had or had not been pre-incubated with 300 nM holo-Tf was used as an analyte. Average kinetic constants and standard errors were computed from triplicate trials.

**Figure S3.**
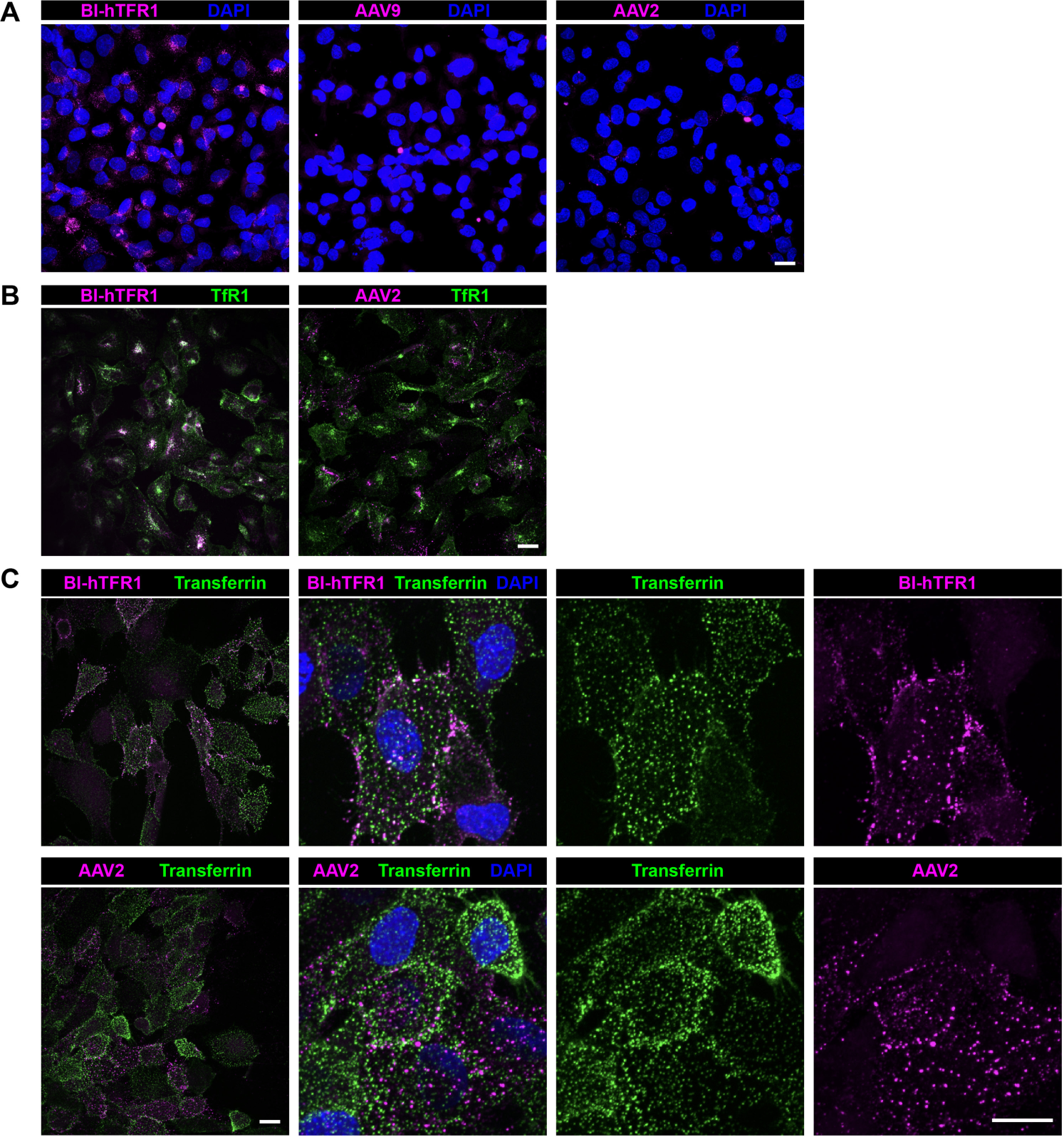
AAV colocalizes with TfR1 and Tf in brain endothelial cells. (**A**) The indicated AAVs were incubated with hCMEC/D3 cells at 25,000 vg/cell for one hour at 37°C. Cells were then fixed, permeabilized, and stained for AAV (magenta) and nuclei (blue). AAV2 was chosen as a control for subsequent experiments because AAV9 immunostaining was nearly undetectable. (**B**) BI-hTFR1 or AAV2 was incubated with hCMEC/D3 cells at 30,000 vg/cell for one hour at 37°C. Cells were fixed, permeabilized, and stained for AAV (magenta) and TfR1 (green). (**C**) BI-hTFR1 or AAV2 was incubated with hCMEC/D3 cells at 25,000 vg/cell in prechilled media containing Tf-647 (1 µg/mL) for one hour at 4°C. Cells were then fixed and stained for AAV (magenta), Tf (green), and nuclei (blue). All scale bars = 15 µm.

**Figure S4.**
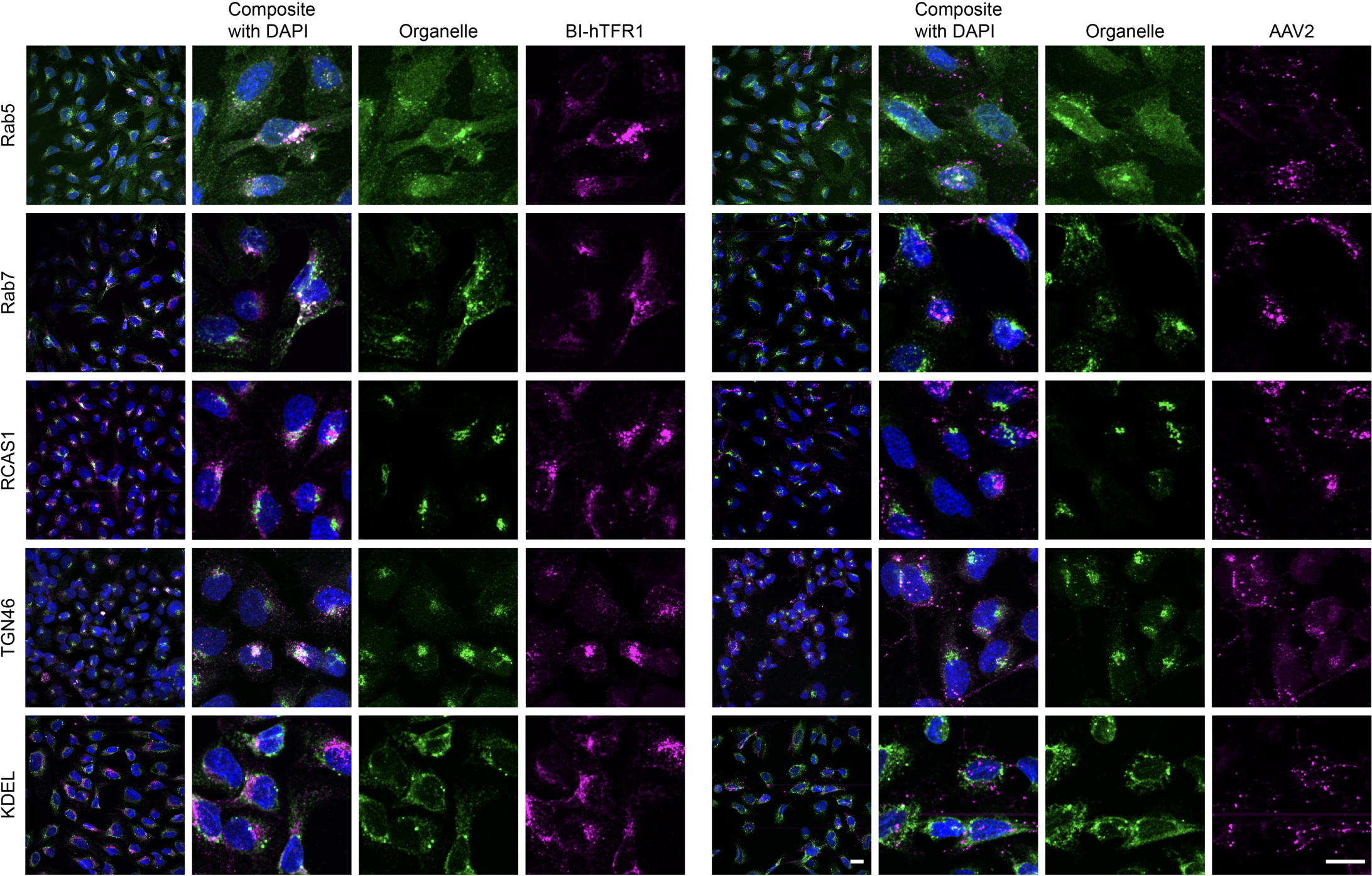
AAV colocalization with subcellular compartment markers. BI-hTFR1 or AAV2 was incubated with hCMEC/D3 cells at 25,000 vg/cell for one hour at 37°C. Cells were then fixed, permeabilized, and stained for AAV (magenta), endosomes (Rab5 and Rab7), cis-Golgi (RCAS1), trans-Golgi network (TGN46) or endoplasmic reticulum (KDEL), and nuclei (blue). All scale bars = 15 µm.

**Figure S5.**
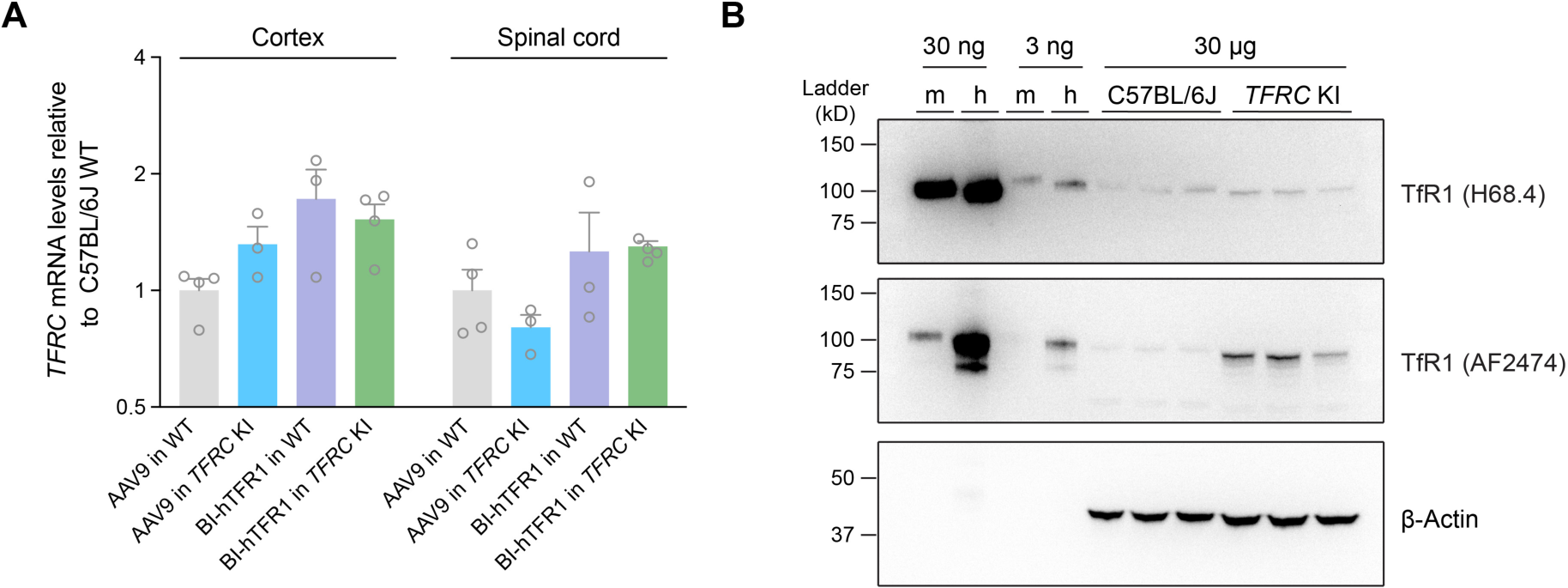
TfR1 levels in *TFRC* KI mice match those in WT mice. (**A**) RT-qPCR using primers spanning exon 1 and exon 2 of the mouse *Tfrc* gene (table S1) was performed to assess mRNA levels in *TFRC* KI mice or C57BL/6J controls treated with either AAV9 or BI-hTFR1 (*n* = 3 or 4 animals per group; error bars indicate ± SEM). No significant difference in TfR1 mRNA between the C57BL/6J and *TFRC* KI mice was detected in the cortex or spinal cord (two-way ANOVA). (**B**) Western blots of protein from the brain tissues of C57BL/6J and *TFRC* KI homozygous mice injected with AAV9 or BI-hTFR1 and purified full-length human or mouse TfR1 Fc-fusion proteins with N-terminal flag tag were stained using a polyclonal anti-human TfR1 antibody, AF2474 (R&D systems), or a monoclonal antibody, H68.4 (Thermofisher, Catalog #13-6890) that recognizes mouse and human TfR1.

**Figure S6.**
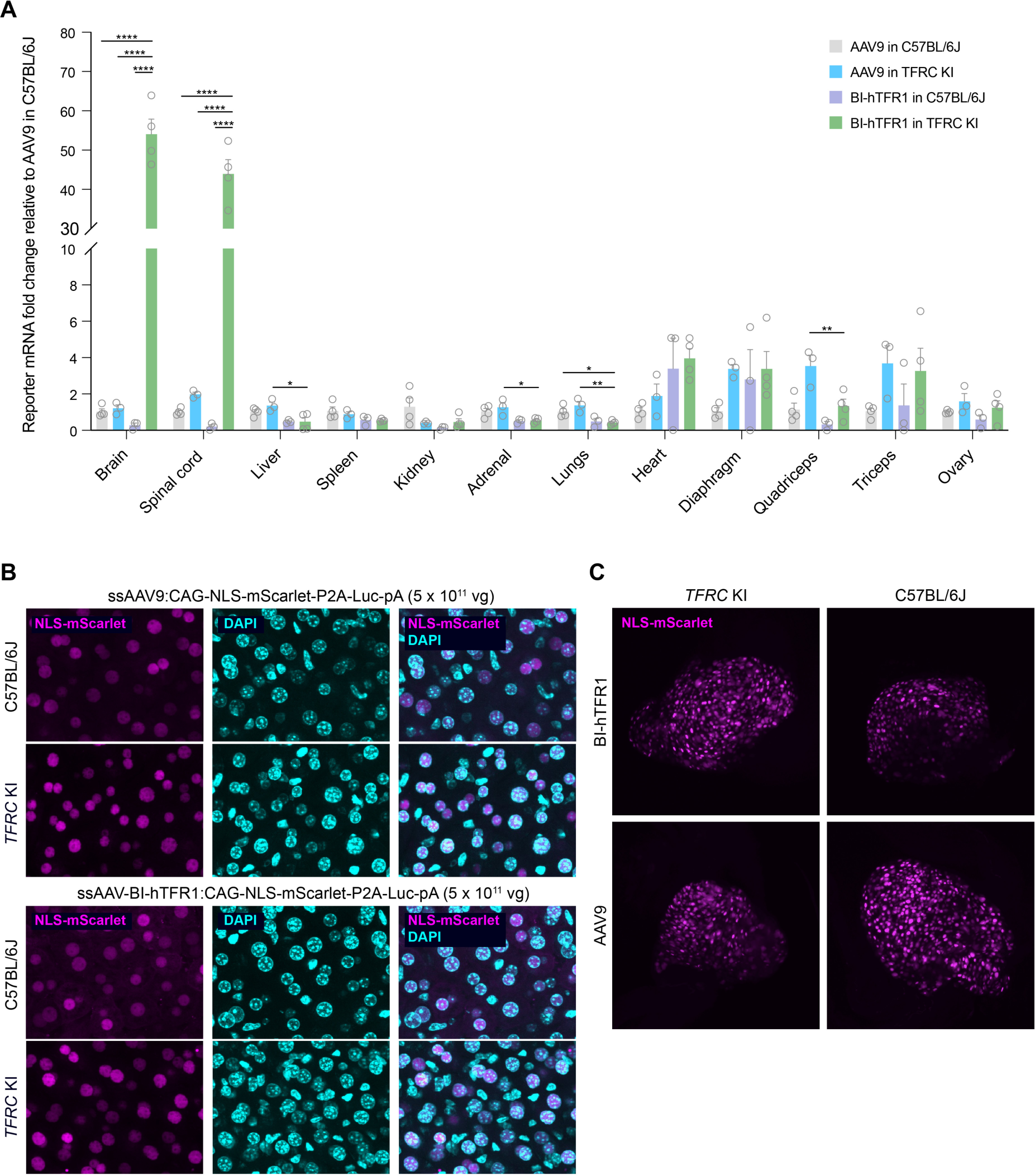
BI-hTFR1 exhibited a CNS-specific enhanced tropism in *TFRC* KI mice and similar transduction to AAV9 in the liver and dorsal root ganglion. BI-hTFR1 or AAV9:CAG-NLS-mScarlet-P2A-Luciferase-WPRE-SV40pA was intravenously injected into adult female C57BL/6J or *TFRC* KI mice at a dose of 5 x 10^11^ vg/mouse. The (**A**) mScarlet transcript levels relative to AAV9 in C57BL/6J mice (two-way ANOVA using BI-hTFR1 in *TFRC* KI mice as the main comparison group with Bonferroni multiple comparison correction: ****, ***, **, and * indicate *p* ≤ 0.0001, ≤ 0.001, ≤ 0.01, and ≤ 0.05, respectively; each data point represents an individual mouse, error bars indicate ± SEM), and native fluorescence in the (**B**) liver and (**C**) dorsal root ganglion at three weeks post-injection are shown.

**Figure S7.**
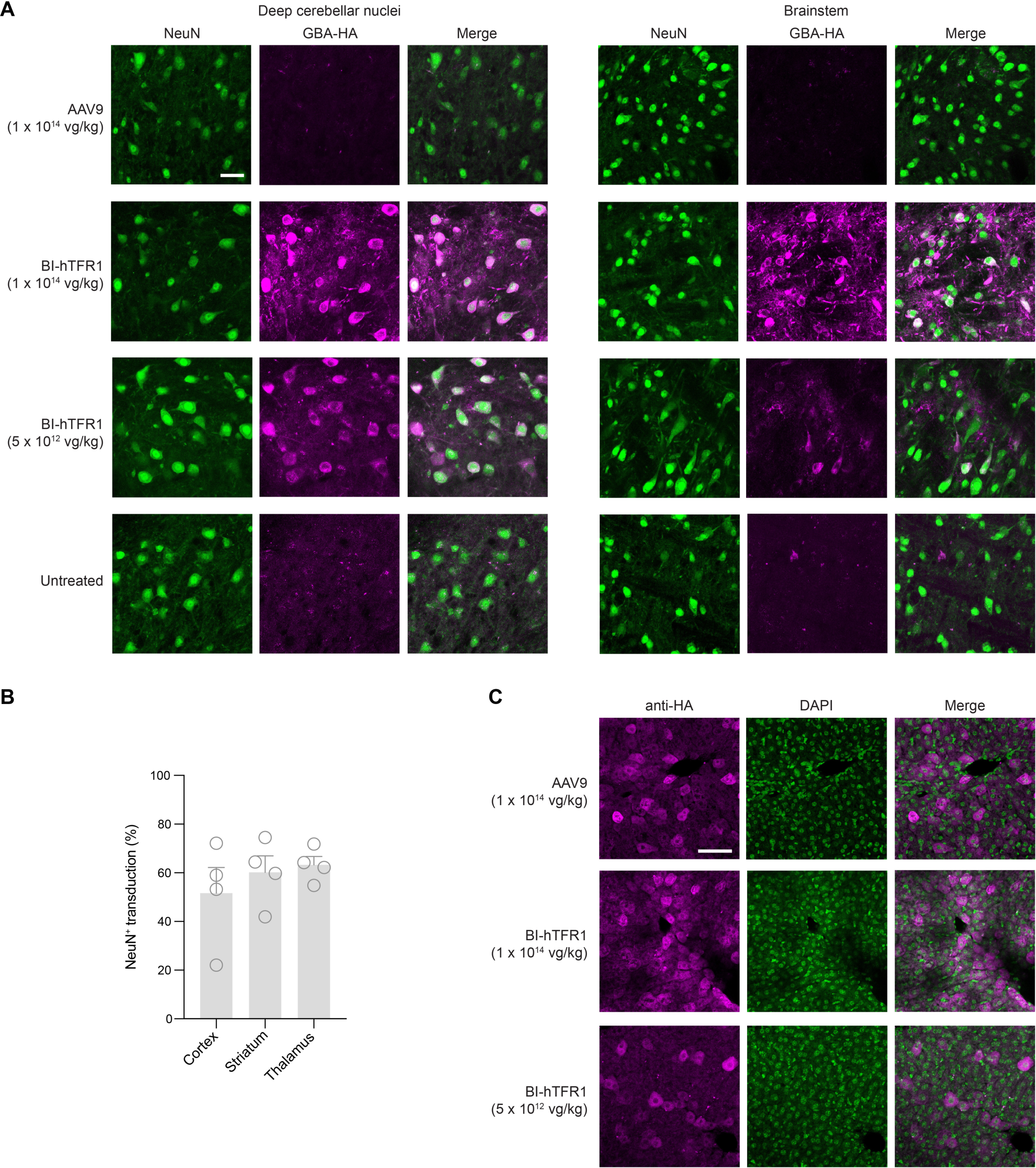
Expression of GBA-HA in the liver and brain regions of mice injected with AAV9 or BI-hTFR1:*GBA1*. (**A**) Images show NeuN (green) and HA (magenta) immunostaining in the deep cerebellar nuclei and brainstem in *TFRC* KI mice with the indicated treatment conditions. Scale bar = 25 µm. (**B**) The cell counts of transduced NeuN+ cells expressing GBA-HA in the cortex, striatum, and thalamus of *TFRC* KI mice injected with 1 x 10^14^ vg/kg BI-hTFR1:*GBA1* are shown. Each data point represents a mouse (*n* = 4). Error bars indicate ± SEM. (**C**) Representative images of transduced liver hepatocytes stained for HA in the indicated conditions are shown. Scale bar = 50 µm.

**Figure S8.**
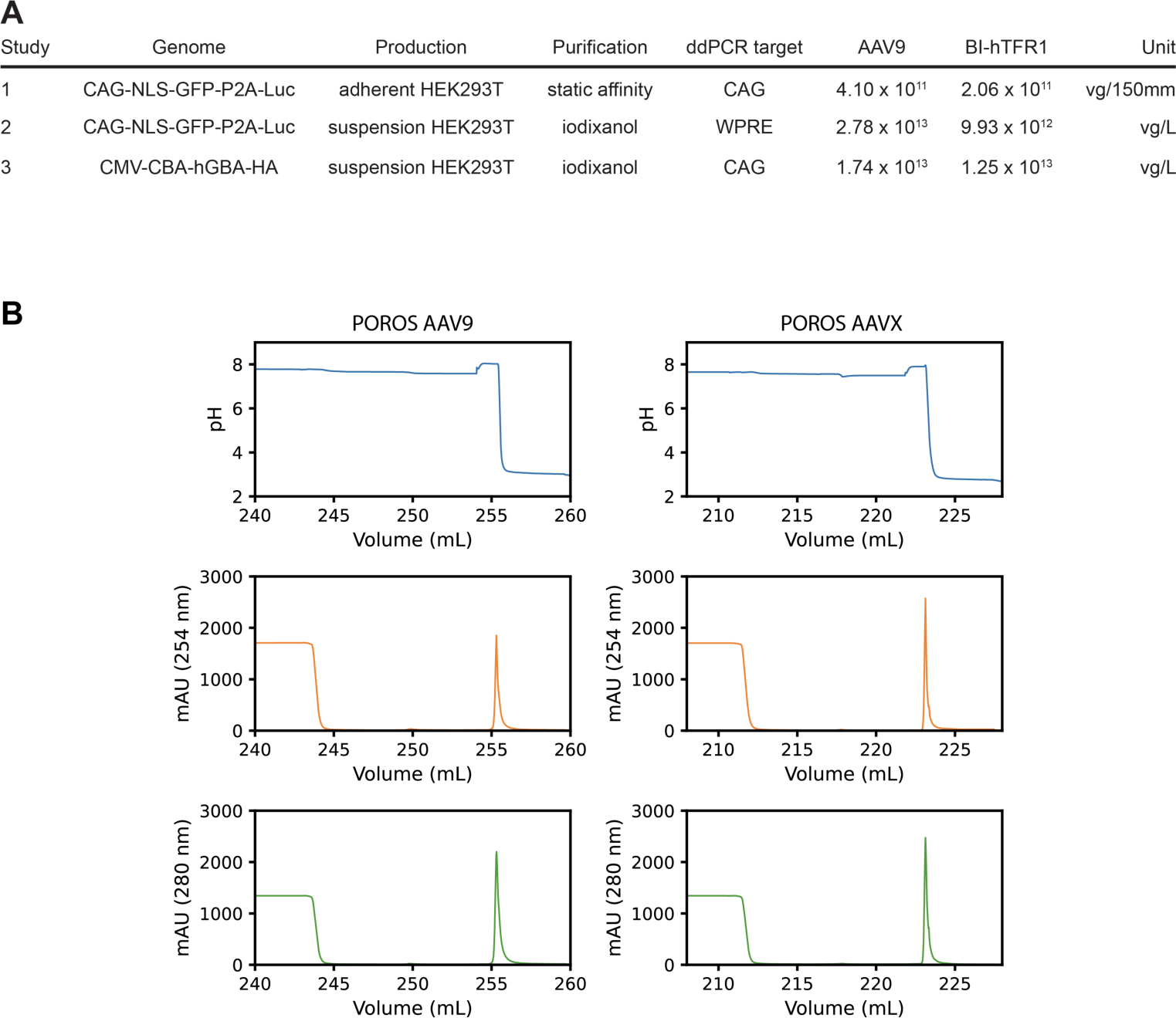
BI-hTFR1 produces comparably to AAV9 and is compatible with purification using POROS CaptureSelect resins. (**A**) In this study, BI-hTFR1 and AAV9 were produced in parallel using either adherent or suspension HEK293T and purified via static affinity or iodixanol. For some of the preparations, the post-purification yields were assessed by ddPCR of either the CAG or WPRE component of the AAV genome and are reported here. (**B**) We separately assessed the compatibility of BI-hTFR1 with affinity purification using POROS CaptureSelect AAV9 and AAVX resin. The top row of chromatograms shows the elution phase indicated by the drop in pH. The second and third rows of chromatograms illustrate the corresponding elution peaks measured by absorbance at 254 nm and 280 nm, respectively. AAV was eluted from POROS CaptureSelect AAV9 using Buffer A at a pH of 3.0 and from POROS CaptureSelect AAVX using Buffer B at a pH of 2.5.

**Table S1.**
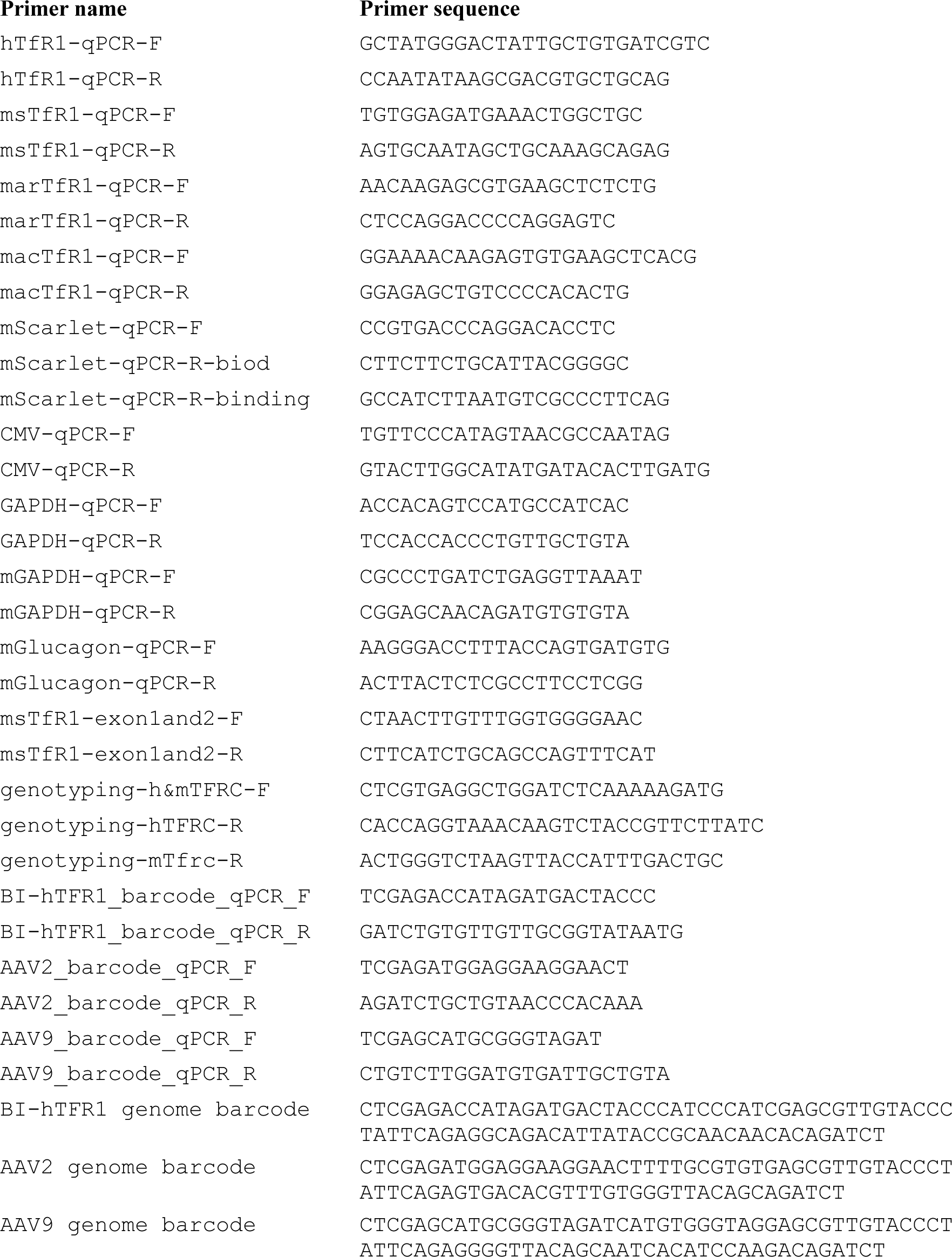
Primers and barcodes used for qPCR in this study. Each sequence is listed 5’ to 3’.

